# Bacterial exo-α-sialidases subvert the complement system through desialylation

**DOI:** 10.64898/2026.02.05.703967

**Authors:** Kurni Kurniyati, Nicholas D. Clark, Qin Fu, Sheng Zhang, Weigang Qiu, Michael G. Malkowski, Chunhao Li

## Abstract

The complement system is a central pillar of innate immunity, yet how bacterial glycan-modifying enzymes subvert complement function remains poorly understood. Bacterial sialidases remove terminal sialic acids from host sialoglycans, but their direct impact on complement immunity is unclear. Here, we investigate the impact of six sialidases from five human pathogens on complement immunity using an integrated approach combining genetics, biochemistry, glycobiology, mass spectrometry, and structural biology. We demonstrate that major complement components (IgG, C1q, C4, and C5) and regulators (Factor I, Factor H, and C4bp) are sialylated, and that bacterial sialidase-mediated desialylation suppresses complement activation and surface deposition, thereby enabling complement evasion. Despite extensive sequence diversity, biochemical and structural analyses reveal that all examined sialidases share a conserved six-bladed β-propeller catalytic domain and cleave both N-acetylneuraminic and N-glycolylneuraminic acids, the predominant mammalian sialic acids. Together, these findings uncover a conserved mechanism by which diverse bacterial pathogens disable complement immunity through desialylation.

## INTRODUCTION

The complement system is a central component of innate immunity, providing a rapid and versatile first line of defense against microbial infections^1^. Complement activation proceeds through the classical, lectin, and alternative pathways, all of which converge on the formation of C3 convertase. Downstream proteolytic cascades amplify immune recognition and effector functions, ultimately driving pathogen opsonization, formation of the membrane attack complex (MAC), and target cell lysis^2,3^. Beyond antimicrobial defense, the complement system also functions as a key regulator of immune homeostasis, bridging innate and adaptive immunity, shaping antibody responses, and facilitating clearance of apoptotic cells^1,4^. Given its potent effector capacity, complement activity is stringently controlled to prevent pathogenic effects, and its dysregulation contributes to a wide spectrum of infectious, inflammatory, and autoimmune diseases^5,6^.

More than 30 plasma proteins constitute the human complement system, the majority of which are glycoproteins primarily bearing N-linked glycans^7^. Glycosylation is essential for the structural stability and function of component factors, including C3, C4, and C5, which require N-linked glycans for proteolytic activation and convertase assembly^7-9^. Perturbation of glycosylation can impair complement function or lead to aberrant activation, e.g., non-glycosylated IgG induces antibody-dependent cellular cytotoxicity; aberrant C3 glycosylation is associated with early onset type 1 diabetes^10,11 912^. Intriguingly, many complement glycoproteins harbor terminal sialic acids, a modification known as sialylation^7,13^. For instance, Factor H (FH), a negative regulator of complement, is glycosylated at nine positions, eight of which carry sialoglycans^14^. C4 is glycosylated at five positions, at least three of which are sialylated^7,15^. IgG is sialylated at Asn297 within its Fc region, a modification required for its anti-inflammatory activity; loss of Fc sialylation abolishes this immunomodulatory function and promotes pathogenic autoantibody responses^16,17^. However, the functional roles of sialylation across most complement factors remain poorly defined.

Sialic acids are a family of nine-carbon sugars typically found at the terminal positions of sialoglycans^18,19^. N-acetylneuraminic acid (Neu5Ac) and N-glycolylneuraminic acid (Neu5Gc), the two most common mammalian sialic acids, play crucial roles in immune recognition and cell-cell interactions^20^. Sialidases (neuraminidases) are a group of structurally diverse enzymes that remove sialic acids from sialoglycans, mainly spanning glycoside hydrolase (GH) families 33, 34, 58, and 83^21-23^. Bacterial pathogens predominantly encode GH33 exo-α-sialidases, a group of enzymes that cleave terminal α-linked sialic acid residues^24^. Many pathogens exploit sialidases to scavenge host sialic acids as nutrient sources or as “molecular disguises” to evade immune recognition^25^. Increasing evidence indicates that bacterial sialidases also promote complement evasion, e.g., the sialidase of *Streptococcus pneumoniae* impairs complement activation, though the precise mechanism is unclear^26^. We recently showed that two oral pathogens, *Porphyromonas gingivalis* and *Treponema denticola*, achieve serum resistance through desialylating key complement factors^27,28^. However, whether this strategy represents a niche-specific adaptation confined to oral pathogens or a broadly conserved mechanism employed by diverse bacterial pathogens across different ecological niches remains unknown.

Here, we address this gap by defining complement sialylation and determining how bacterial sialidases subvert complement immunity. We first characterized the complement sialylation using glycosylation staining, lectin blotting, and mass spectrometry. Using NanH from *T. denticola* as a model of GH33 sialidases, we demonstrate that desialylation of complement factors directly impairs complement-mediated killing and then elucidate its underlying catalytic mechanism through functional and structural analyses. Finally, we show that this complement-subverting mechanism is conserved across six representative sialidases from five human pathogens, including *Bacteroides fragilis, Clostridium perfringens, Tannerella forsythia, T. denticola*, and *Vibrio cholerae*. Together, our findings reveal a conserved glycan-targeted strategy of complement subversion that transcends phylogenetic and ecological boundaries, positioning bacterial sialidases as key virulence determinants and potential targets for the development of novel antimicrobial modalities.

## RESULTS

### Human serum proteins including complement factors are extensively sialylated

Human serum contains a broad repertoire of proteins, many of which are glycosylated^7,29,30^. To determine if human serum contains sialylated proteins, 2% normal human serum (NHS) was subjected to lectin blots probed against *Sambucus nigra* agglutinin (SNA) and *Maackia amurensis* II (MAL II), two lectins that specifically recognize α2,6-linked and α2,3-linked sialic acids^31,32^, respectively. Concanavalin A (Con A), a lectin that interacts with diverse receptors containing mannose carbohydrates^33^, was used as a control. Multiple bands were detected by these three lectins (**Supplementary Fig. S1**), indicating that human serum is highly enriched in sialylated glycoproteins.

Several complement factors are present at particularly high concentrations in human serum, such as C3 (1,500□µg/ml), FH (500□µg/ml), C4 (315□µg/ml), and C4-binding protein (C4bp, 250□µg/ml)^7^. To determine if these complement factors are modified by sialoglycans, purified IgG, C1q, C4, C5, FH, Factor I (FI) and C4bp (**Fig. 1a**) were selected and subjected to glycosylation staining and lectin blots with SNA and MAL II. All tested complement proteins are positive in glycosylation staining (**Fig. 1b, c**), indicating that they are glycosylated. These proteins also bound to SNA (**Fig. 1d**), suggesting the presence of α2,6-linked sialic acids, particularly on C4bp α-chain, FI, FH, C4 α-chain, C5 α-chain, C1q subunits A and B, and the IgG heavy chain. In contrast, only a subset of complement proteins (IgG, C1q and C4) were recognized by MAL II (**Fig. 1e**), indicating the presence of α2,3-linked sialic acids.

**Figure 1.**
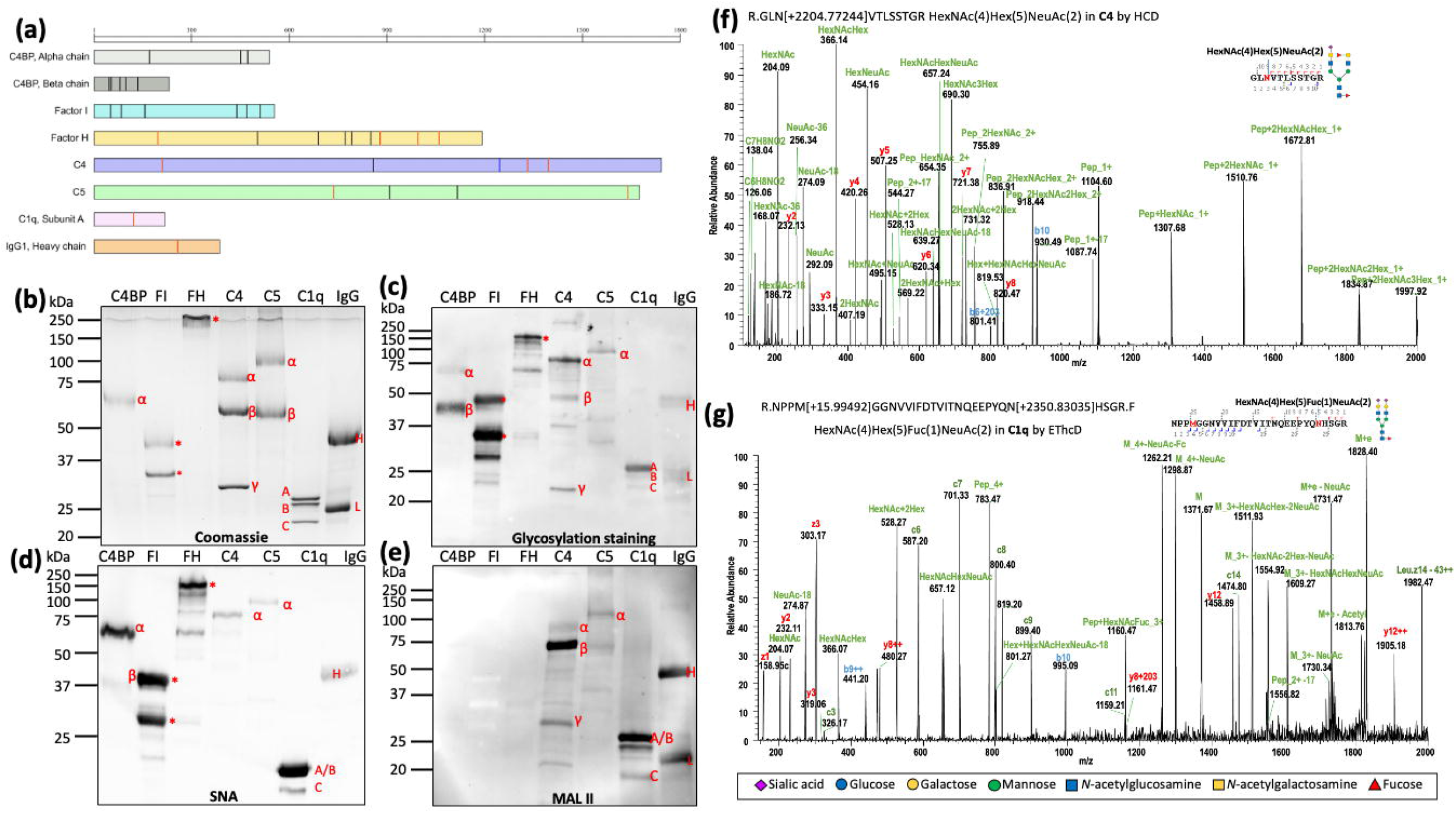
Human serum and complement proteins are extensively glycosylated and sialylated. **(a)** Schematic representation of predicted glycosylation sites in human serum proteins, including complement components C4b-binding protein (C4BP) α and β chains, Factor I (FI), Factor H (FH), C4, C5, C1q subunits A, and the IgG1 heavy chain. Numbers above each schematic indicate amino acid positions starting from the N-terminus following signal peptide cleavage. Glycosylation sites were predicted using UniProt and GlyConnect. Black lines denote predicted N-linked glycosylation sites, blue lines denote predicted O-linked glycosylation sites, and red lines indicate N-linked glycosylation sites experimentally identified in this study. **(b-e)** Complement proteins C4BP, FI, FH, C4, C5, C1q, and IgG were analyzed by SDS–PAGE followed by Coomassie Brilliant Blue staining (b), total glycoprotein staining (c), and lectin blotting using *Sambucus nigra agglutinin* (SNA) (d) and *Maackia amurensis* lectin II (MAL II) (e). Abbreviations: α, alpha chain; β, beta chain; γ, gamma chain; A, subunit A; B, subunit B; C, subunit C; H, heavy chain; L, light chain; ^*^, protein fragments. **(f-g)** Representative glycopeptides identified from C4 (f) and C1q (g) by nanoLC–ESI–MS/MS analysis. Glycan compositions were predicted using GlyConnect and illustrated using DrawGlycan-SNFG.

To confirm the lectin blot results, IgG, C1q and C4 were selected and subjected to nano LC-ESI/ MS/MS analysis. The results showed that they are all modified by N-linked glycans and sialoglycans at multiple sites (**Fig. 1f, g** and **Supplementary Fig. 2 and Table 2-4**). In addition to previously reported sialoglycans^7,14-16,34,35^, several new forms were detected, although their structures remain unresolved. These results collectively demonstrate that human serum, including key complement proteins, is rich in N-linked glycoproteins that are extensively modified with both α2,6- and α2,3-linked sialic acids, highlighting the complexity of their glycosylation profiles.

### GH33 sialidases are diverse but share conserved catalytic features and structural folds

Bacterial sialidases predominantly belong to the GH33 family^24,25^. Phylogenetic analysis of 289 GH33 sequences downloaded from UniProt revealed extensive diversity, comprising 162 tips across 55 bacterial genera (**Fig. 2a**). Most sequences cluster by taxonomy, indicating an ancient evolutionary origin, orthologous diversification, and little evidence of horizontal gene transfer. Sequence alignment of six representative sialidases from five different pathogens (**Fig. 2b**), including *B. fragilis, C. perfringens, T. denticola, T. forsythia*, and *V. cholerae*, showed divergent N-terminal architectures while their C-termini all contain a conserved exo-α-sialidase catalytic domain consisting of a F/YRIP motif and multiple Asp-boxes (**Fig. 2c and Supplementary Fig. 3**). We then compared their crystal structures (PDB) or AlphaFold models (Uniprot entry) using *Td*-NanH, a sialidase of *T. denticola* (TDE0471), as a template, and found that their C-terminal domains all possess conserved exo-α-sialidase active sites (**Supplementary Fig. 4**) and six-bladed β-propeller structures (**Fig. 2d-h**) with C_⍰_ RMSD values ranging from 1.673□ (*Cp*-NanI) to 2.044□ (*Vc-*NanH). Notably, the higher C_⍰_ RMSD for the *Vc-*NanH alignment is likely attributed to the insertion of a large CBM_nc_ domain into the sialidase domain which is absent in the other five sialidases. Together, these analyses demonstrate that bacterial GH33 sialidases possess diverse domain architectures but share a conserved catalytic core and structural fold.

**Figure 2.**
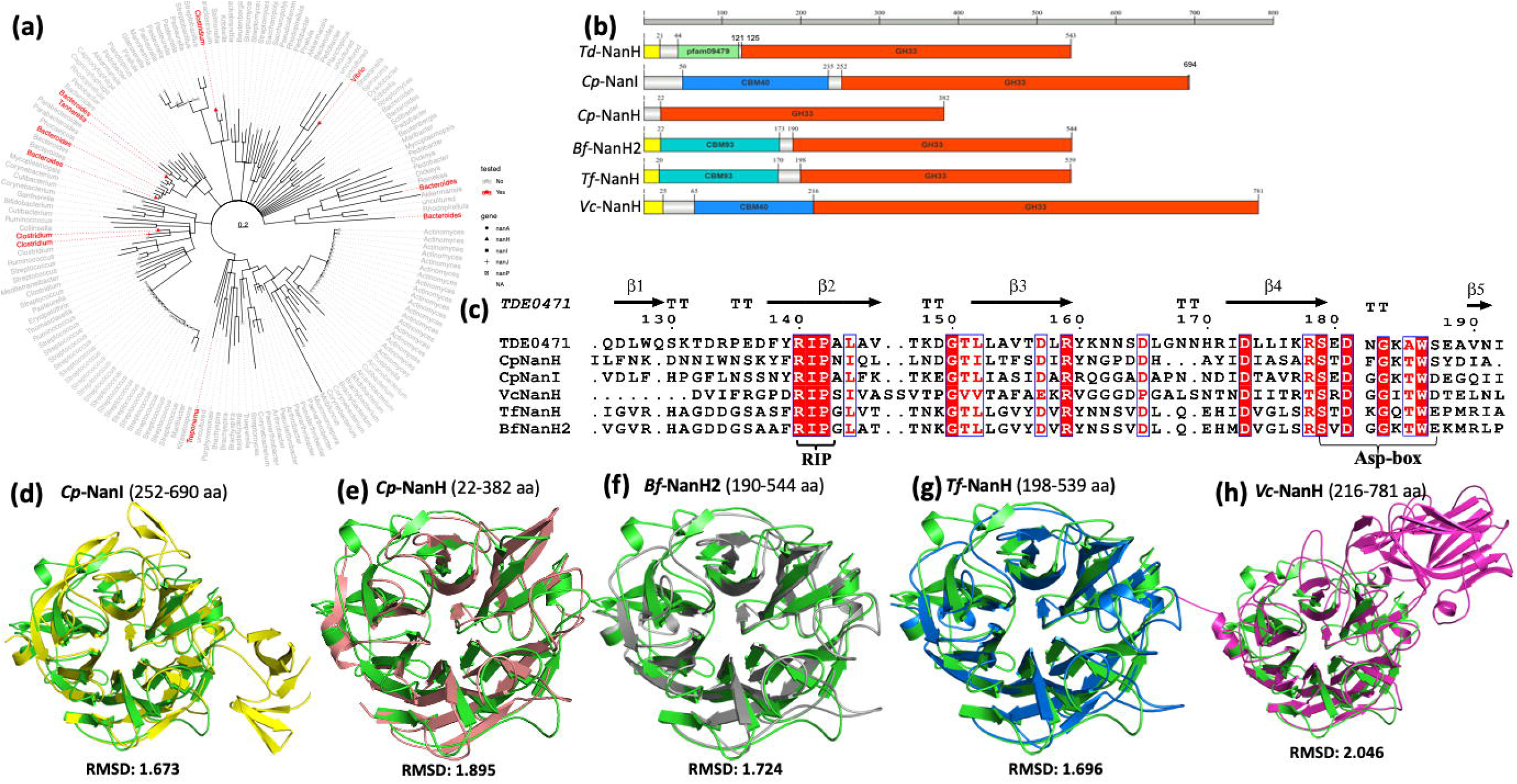
GH33 sialidases are evolutionarily diverse but share conserved catalytic motifs and structural folds. **(a)** Maximum-likelihood phylogenetic tree of bacterial GH33 exo-sialidases. A total of 289 sequences from UniProt and GenBank were aligned using MUSCLE, and a FastTree-based phylogeny was consolidated to 162 representative sequences spanning 55 bacterial genera. The scale bar indicates 0.2 amino acid substitutions per site. Experimentally characterized sialidases are highlighted in red: *Treponema denticola* NanH (*Td*-NanH), *Clostridium perfringens* NanI (*Cp*-NanI) and NanH (*Cp*-NanH), *Bacteroides fragilis* NanH2 (*Bf*-NanH2), *Tannerella forsythia* NanH (*Tf*-NanH), and *Vibrio cholerae* NanH (*Vc*-NanH). Tip symbols indicate distinct gene annotations. **(b)** Domain organization of the six sialidases. Amino acid positions are shown relative to the N-terminus. Domains were annotated using NCBI and CAZy resources. Signal peptides (SignalP-6.0) are shown in yellow, GH33 catalytic domains in red, and additional carbohydrate-binding modules (CBMs) or the pfam09479 domain in other colors. **(c)** Multiple sequence alignment of the conserved catalytic region containing the “Y/FRIP” motif. Secondary structure elements derived from *Td*-NanH are shown above the alignment, with the conserved “Y/FRIP” motif and Asp-box sequences highlighted. **(d–h)** Structural superposition of GH33 sialidases using *Td*-NanH (green) as the reference, overlaid with *Cp*-NanI (yellow; PDB 5TSP), *Cp*-NanH (salmon; PDB 8UB5), *Bf*-NanH2 (gray; AlphaFold), *Tf*-NanH (marine; PDB 7QYP), and *Vc*-NanH (magenta; PDB 1KIT).

### GH33 sialidases desialylate human serum and complement proteins

To assess their enzymatic properties, we expressed and purified six recombinant GH33 sialidases (**Fig. 3a**), including *Bf*-NanH2, *Cp*-NanI, *Cp*-NanH, *Td*-NanH, *Tf*-NanH, and *Vc*-NanH, and then conducted kinetic analyses using 4-methylumbelliferyl-α-D-N-acetylneuraminic acid (4-MUNANA), a widely used fluorogenic substrate for measuring sialidase activity^23,27^. All six enzymes exhibited sialidase activity, albeit with substantial variation in their catalytic efficiency (**Fig. 3b** and **Supplementary Table 5**). Among them, *Td-*NanH exhibited the highest reaction rate (V_max_ = 995.2 ± 25.06 µM/min) but low substrate binding affinity (K_m_ = 840.9 ± 41.70 µM), whereas *Tf-*NanH showed the lowest reaction rate (V_max_ = 96.89 ± 2.61 µM/min) yet the highest binding affinity (K_m_ of 171.9 ± 15.44 µM). These findings indicate that GH33 sialidases are catalytically active yet display markedly variable substrate affinities and turnover rates, suggesting their functional diversity in glycan remodeling potential.

**Figure 3.**
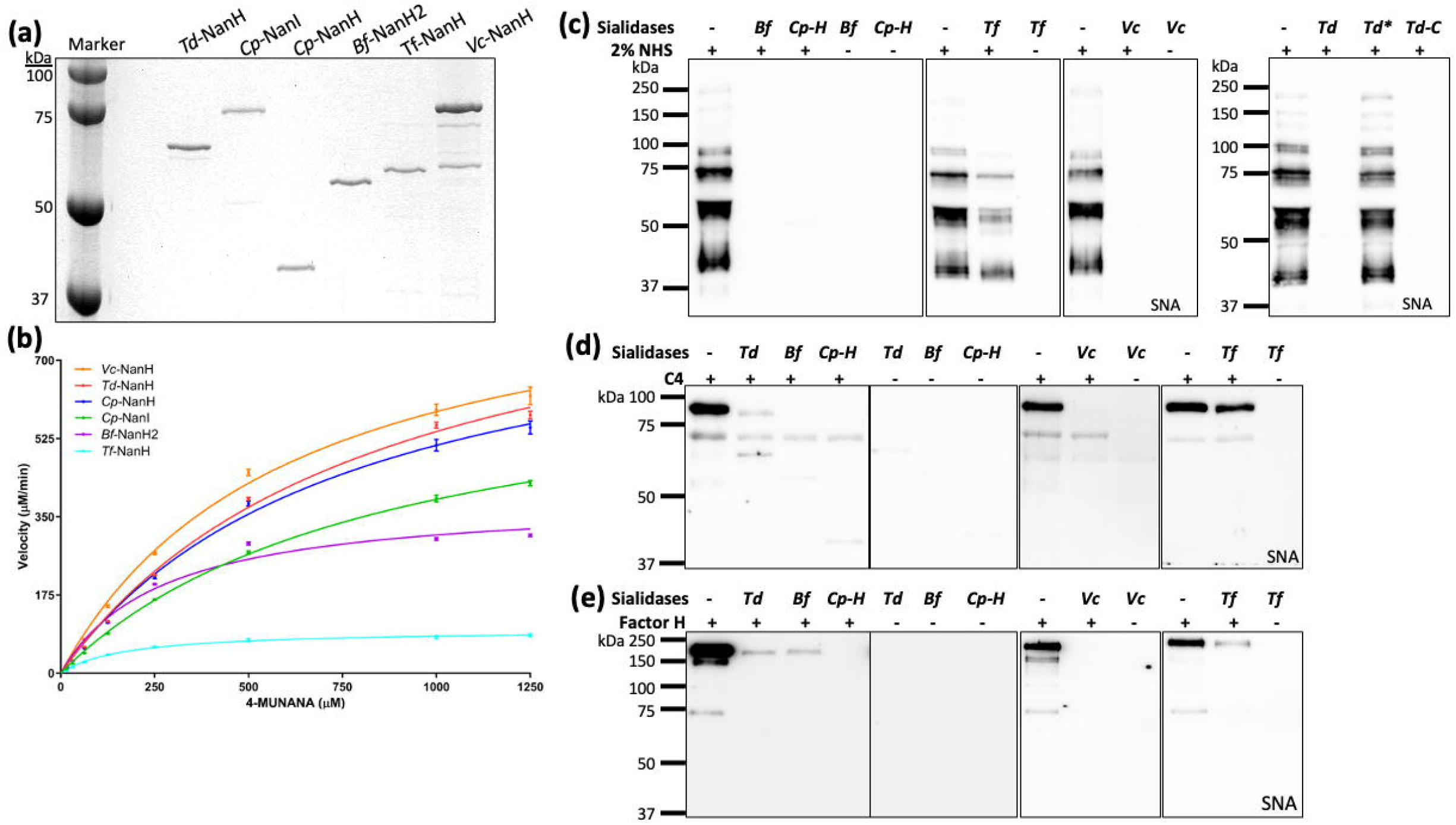
GH33 sialidases desialylate human serum and complement proteins. **(a)** SDS-PAGE analysis of six purified recombinant sialidases, including *Td*-NanH, *Cp*-NanI, *Cp*-NanH, *Bf*-NanH2, *Tf*-NanH, and *Vc*-NanH. *Cp*-NanI was purchased from Sigma-Aldrich, N2133 and the remaining five sialidases were expressed in *E. coli* and purified using FPLC. **(b)** Kinetic analysis of six GH33 sialidases. This assay was carried out as described in the Methods section using 4-methylumbelliferyl-α-D-N-acetylneuraminic acid (4-MUNANA) as a substrate. Saturation curves were fitted to Michaelis-Menten kinetics using GraphPad Prism ; and K_m_ and V_max_ were calculated (see details in Supplementary Table 5). **(c-e)** GH33 sialidases desialylate human serum, C4 and FH proteins. For this study, normal human serum (NHS), C4 or FH proteins were incubated with six recombinant sialidases at 37°C for 1 or 3 hours, followed by SDS-PAGE and lectin blots using SNA. Abbreviations: Td, *Td-*NanH; Cp-I, *Cp-* NanI; Cp-H, *Cp-*NanH; Bf, *Bf-*NanH2; Tf, *Tf-*NanH; and Vc, *Vc-*NanH.

We next evaluated the ability of these enzymes to desialylate human serum proteins. Normal human serum (NHS, 2%) was incubated with recombinant sialidases at 37 °C for 1 or 3 h, followed by lectin blot analysis using SNA (**Fig. 3c**). Five sialidases markedly reduced SNA signals after 1 h, whereas *Tf*-NanH, consistent with its lower catalytic activity, exhibited weaker desialylation, with residual SNA reactivity detectable even after 3 h incubation. Similar results were observed when the analysis was performed using MAL II lectin blot (**Supplementary Fig. 5**). Comparable desialylation activity was observed when purified complement proteins C4 and factor H (FH) were treated with the recombinant enzymes (**Fig. 3d, e** and **Supplementary Fig. 6 & 7**). Notably, the catalytically inactive mutant *Td*-NanH^*^ (RIP motif mutated) failed to remove sialic acids from these substrates, confirming that enzymatic activity is required for desialylation. Collectively, these results demonstrate that GH33 sialidases are active glycosidases capable of removing terminal sialic acids from human serum and complement proteins.

### Sialidase-based mapping of complement protein sialylation

Several complement factors (e.g., C1q and FH) are modified by sialoglycans^7,14,17,34^; however, their precise sites and linkages (e.g., α2,3 versus α2,6) remain poorly defined. To delineate their sialylation patterns, human serum and seven complement proteins (C1q, C4, C5, C4bp, FH, FI, and IgG) were treated with recombinant *Td*-NanH, a representative sialidase from a panel of six enzymes. SNA and MAL II lectin blots showed that *Td*-NanH removed sialic acids from human serum in a dose-dependent manner (**Fig. 4a-c and Supplementary Fig. 8**). Lectin blot analyses revealed that SNA reactivity in C4bp, FI, FH, C4, and C5 was completely abolished following 3 h *Td*-NanH treatment (**Fig. 4d and Supplementary Fig. 9 & 10**), indicating predominant α2,6-linked sialylation. In contrast, C1q and IgG were resistant to *Td*-NanH-mediated desialylation, suggesting that they are modified by alternative sialic acid linkages such as α2,3.

**Figure 4.**
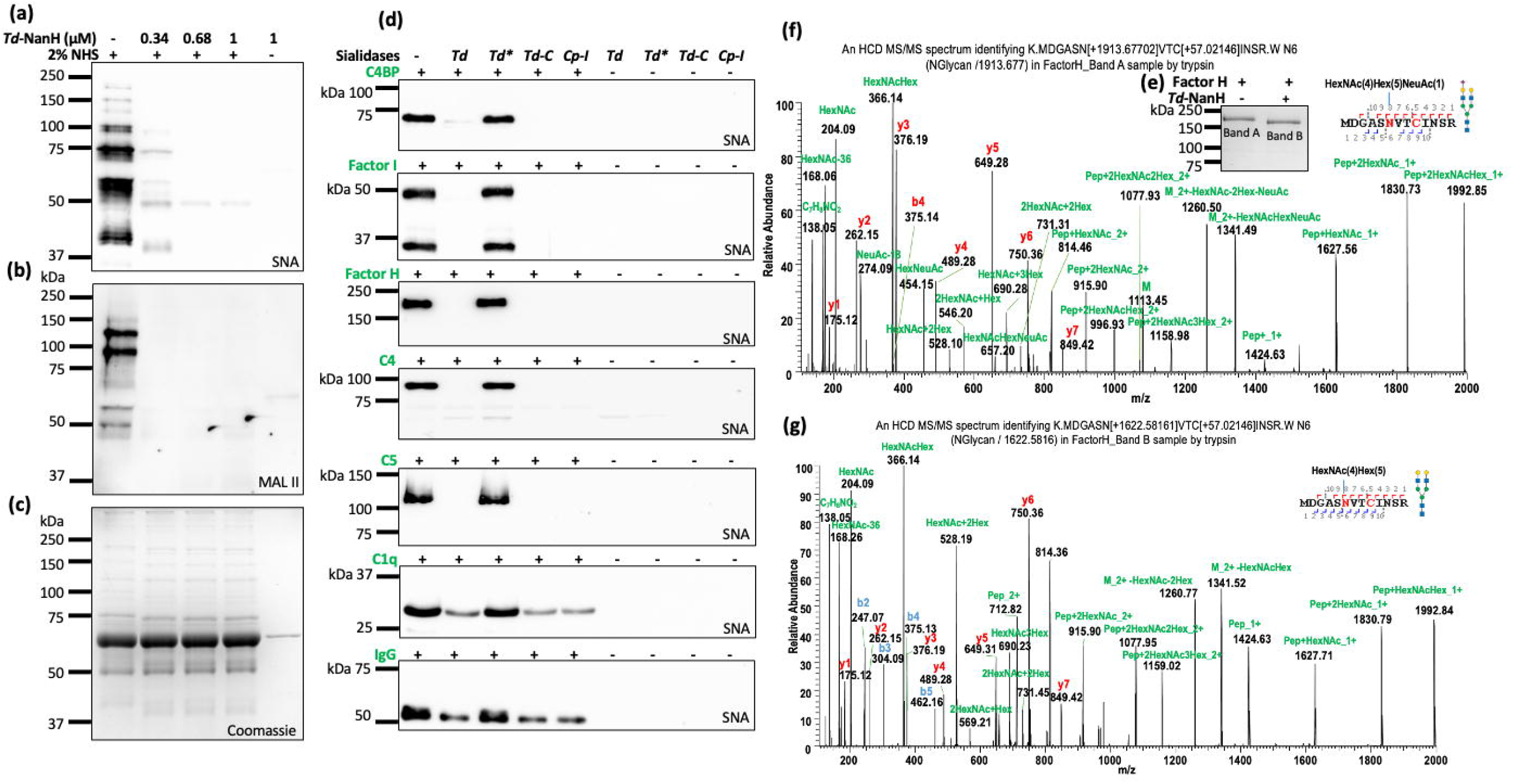
*Td*-NanH desialylates human serum and complement proteins. **(a-c)** *Td*-NanH removes human serum sialic acids in a dose dependent manner. Normal human serum (NHS) was treated with varying amount of *Td-* NanH at 37°C for 1 or 3 hours. Samples were analyzed by SDS-PAGE, followed by lectin blot analysis using SNA (a) and MAL II (b) and Coomassie staining (c). **(d)** *Td*-NanH desialylates seven complement proteins. For this study, seven complement proteins, including C4BP, FI, FH, C4, C5, C1q, and IgG, were incubated sialidases at 37°C for 1 or 3 hours and then subjected to SDS-PAGE followed by lectin blots using SNA. *Td*^*^: a catalytically inactive mutant of *Td*-NanH (R140A P142A); *Td*-C: the C-terminal fragment of *Td*-NanH. *Cp*-I was included as a control. **(e-g)** Mapping the glycosylation sites of Factor H (FH). For this study, FH was incubated with or without *Td*-NanH for 3 hours at 37□°C and then subjected to SDS-PAGE (e). Untreated (Band A) and treated (Band B) samples were excised for nano LC-ESI/MS/MS analysis. (f, g) A representative nanoLC-ESI-MS/MS spectrum of a C5 glycopeptide detected from untreated and treated FH. The corresponding glycans were predicted using GlyConnect and illustrated using DrawGlycan-SNFG.

To define sialylation sites, treated and untreated FH and C5 were analyzed by nano LC-ESI-MS/MS (**Fig. 4e-g**). In untreated FH, ten sialoglycans were identified at Asn882, Asn911, and Asn1029 with 49% sequence coverage, whereas *Td*-NanH treatment resulted in near-complete loss of sialoglycans despite increased glycan coverage (52%) (**Fig. 4e-g** and **Supplementary Table 7**). A similar pattern was observed for C5 (**Supplementary Fig. 11 and Table 6**). Together, these data confirm that multiple complement proteins are sialylated and demonstrate that sialidase treatment combined with lectin blotting and mass spectrometry provides a robust strategy for precise mapping of sialylation sites and linkage types on complement proteins.

### Sialidase-mediated desialylation impairs serum killing

Among the six sialidases tested, *Td-*NanH, a surface-exposed sialidase of *T. denticola*^28^, exhibited the highest catalytic efficiency and desialylation activity toward human serum proteins and multiple complement factors (**Fig. 4a-d**). Accordingly, *Td*-NanH was selected as a representative enzyme to evaluate the impact of sialidase-mediated desialylation on complement activation and serum bactericidal activity. *Td*-NanH contains a conserved ^140^RIP motif and three Asp-boxes characteristic of GH33 sialidases (**Fig. 5a**). Kinetic analyses demonstrated that *Td*-NanH hydrolyzed both 4-MUNANA and 4-methylumbelliferyl-α-D-glycolylneuraminic acid (4-MUNAGA), which represent Neu5Ac and Neu5Gc, respectively, albeit with distinct reaction rates (**Fig. 5b, c**). Site-directed mutagenesis of residues R140 and P142 (*Td*-NanH^*^) completely abolished its sialidase activity (**Fig. 5b**), confirming their essential role in catalysis.

**Figure 5.**
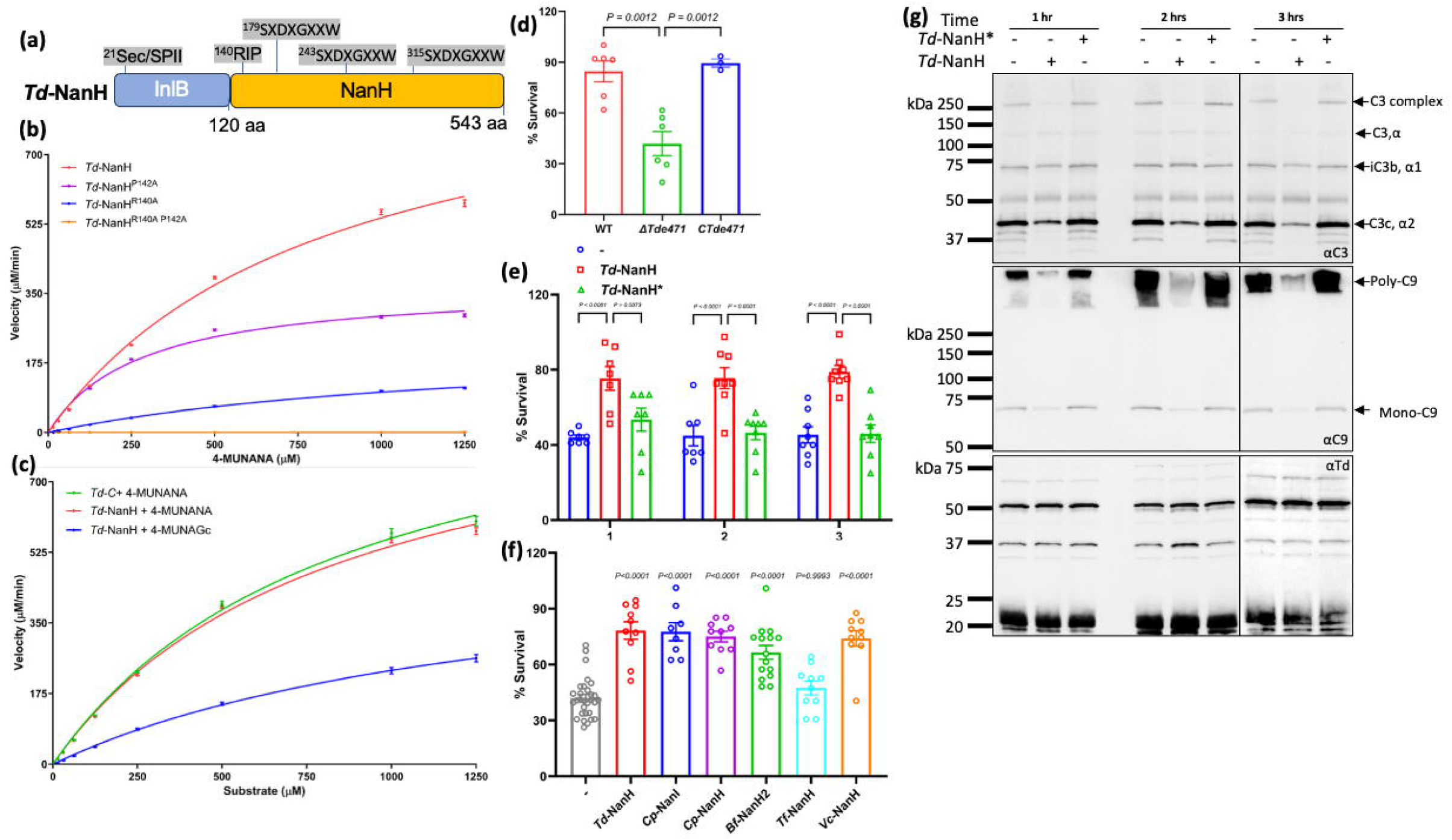
Sialidases protect bacteria from serum-mediated killing. **(a)** Schematic illustration of the domain architecture of *T. denticola* NanH (*Td*-NanH). Annotated features include the signal peptide, the conserved “RIP” motif, and three Asp-box motifs with their respective amino acid positions. InlB: *Listeria monocytogenes* Internalin B. **(b-c)** Enzymatic kinetics of *Td*-NanH toward the fluorogenic substrates 4-MUNANA and 4-MUNAGc. Assays were performed as described in Figure 3 using wild-type *Td*-NanH and three point mutants including *Td*-NanH^R140A^, *Td*-NanH^P142A^, and *Td*-NanH^R140AP142A^; and as C-terminal fragment of *Td*-NanH (*Td-C*) **(d)** Serum bactericidal assays. *T. denticola* wild-type, the *Td*-NanH deletion mutant (Δ*Tde471*), and the isogenic complemented strain (*CTde471*) were incubated with 25% normal human serum (NHS) or heat-inactivated serum (HIS) for 30 min at 37 °C. Viable bacteria were quantified using a Petroff– Hausser counting chamber. Survival was calculated as the ratio of viable cells in NHS relative to HIS. Data are presented as mean ± SEM and analyzed by one-way ANOVA with Tukey’s multiple comparisons test (*P < 0*.*01*). **(e)** *Td*-NanH restores serum resistance in the Δ*Tde471* mutant. NHS (25%) was pre-treated with recombinant *Td*-NanH or the catalytically inactive mutant *Td*-NanH^*^ for 1-3 h at 37 °C prior to serum killing assays with the Δ*Tde471* strain using the same protocol described in (d). **(f)** GH33 sialidases broadly inhibit serum bactericidal activity. NHS (25%) was pre-treated with six GH33 sialidases for 1 h at 37 °C and subsequently used in serum killing assays with the Δ*Tde471* mutant, following the protocol in (d). **(g)** Complement deposition on Δ*Tde471* cells. A total of 5 × 10□ Δ*Tde471* cells were incubated with 25% NHS for 15 min at 37 °C under three conditions: untreated NHS, *Td*-NanH-treated NHS, or *Td*-NanH^*^-treated NHS. Cell-associated complement components were analyzed by SDS-PAGE and immunoblotting using antibodies against C3 and C9. A *T. denticola* specific antibody served as a loading control.

Loss-of-function analyses revealed that *Td*-NanH contributes to serum resistance in *T. denticola*. Deletion of the gene (*TDE0471*) encoding *Td*-NanH significantly reduced bacterial survival in 25% normal human serum (**Fig. 5d**), whereas pretreatment of serum with recombinant *Td*-NanH restored bacterial survival (**Fig. 5e**). Similar rescue experiments using the other five recombinant sialidases showed varying degrees of protection against serum killing (**Fig. 5f**), with survival rates ranked as follows: *Cp*-NanI (78%), *Cp*-NanH (75%), *Vc*-NanH (73%), *Bf*-NanH2 (67%), and *Tf*-NanH (47%). We also measured the impact of *Td*-NanH on complement activation and deposition and found that pretreatment of human serum with active *Td*-NanH, but not the inactive *Td-*NanH^*^ mutant, significantly reduced complement activation and deposition on the surface of *T. denticola* cells (**Fig. 5g**). Collectively, these results demonstrate that GH33 sialidases protect bacteria from complement-mediated killing by desialylating serum complement factors.

### Structural analyses of unliganded *Td*-NanH

To elucidate the catalytic mechanism of GH33 sialidases, we first determined the crystal structure of unliganded *Td*-NanH at 1.63 Å resolution (**Fig. 6**). *Td*-NanH comprises two major domains: an N-terminal domain (N0471; residues 44-121) with an undefined function and a C-terminal catalytic domain (C0471; residues 122-543) adopting the canonical six-bladed β-propeller fold characteristic of GH33 sialidases (**Fig. 6a**). The N0471 domain is comprised of one anti-parallel β-sheet and one half-turn of an ⍰-helix, forming a β-grasp fold. The β-sheet is formed by four individual β-strands which form a concave surface that faces the C0471 domain. The concave surface is filled by the strand connecting β1 to β2 which contains the half-turn helix, occluding the surface. On the N-terminal side of the β-sheet, a second, highly electronegative concave surface is formed. We observed additional electron density with this surface which we subsequently modeled as a bis-tris propane molecule, citrate molecule and two cadmium ions (**Supplementary Fig.12**), derived from the crystallization cocktail. Structural comparison revealed that N0471 is structurally similar to Internalin B (InlB) from *Listeria monocytogenes* (**Fig.6b-d**)^36,37^.

**Figure 6.**
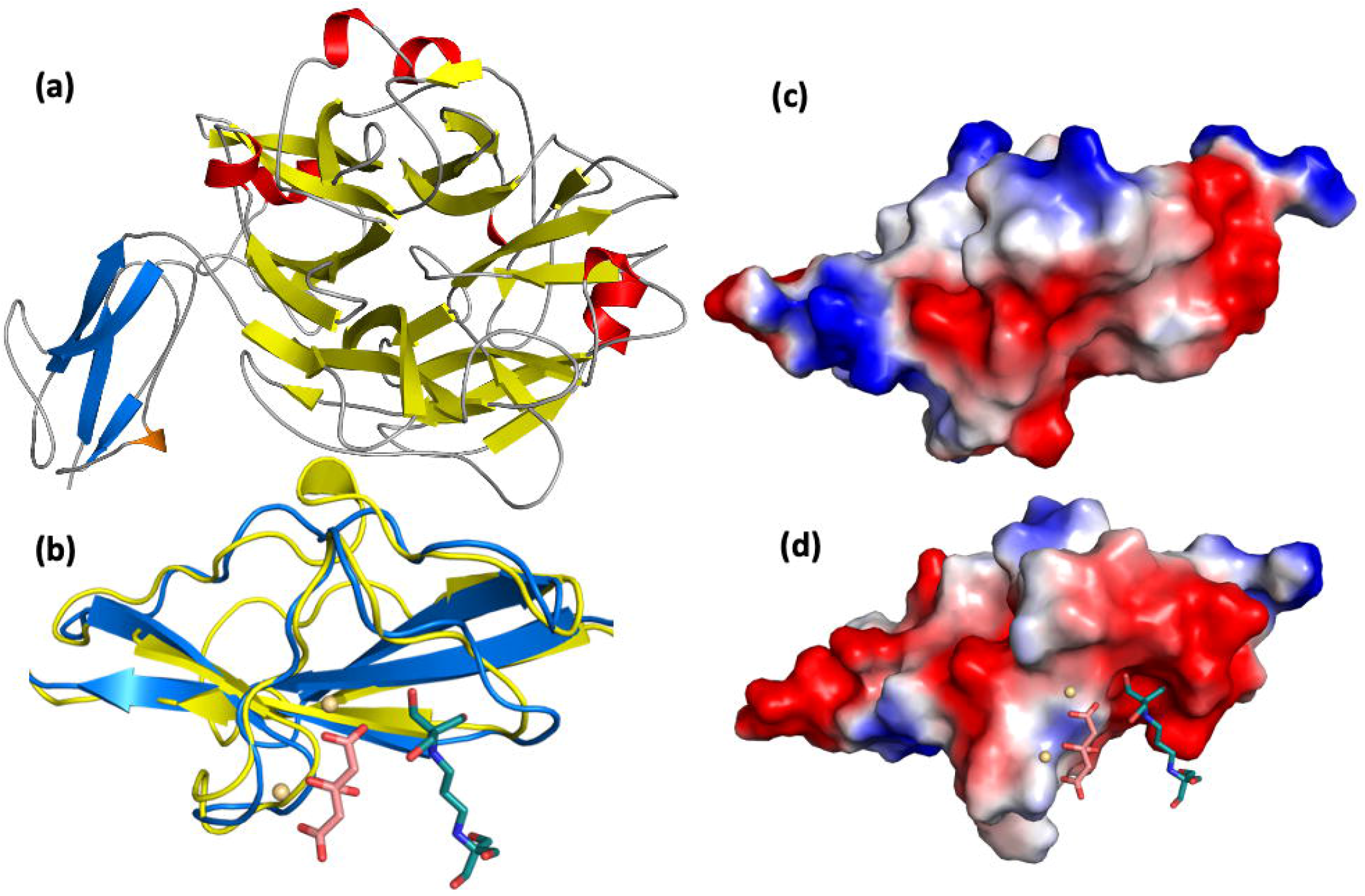
Global architecture of *Td*-NanH. **(a)** Cartoon representation of the global structure *Td*-NanH highlighting the two domains. The N-terminal domain (N0471) is shown with the β-sheets in blue and the ⍰-helix in orange. The canonical C-terminal sialidase domain (C0471) is shown with the β-sheets and ⍰-helices in yellow and red, respectively. The active site is located at the center of the β-propeller domain in C0471. **(b)** Structural overlay of N0471 and InlB. InlB and N0471 are shown in cartoon representation in blue and yellow, respectively. Citrate molecule, bis-tris propane molecule, and cadmium ions captured in N0471 from the crystallization cocktail are shown as salmon sticks, dark teal sticks, and wheat spheres, respectively. **(c-d)** Electrostatic surface maps of InlB (c) and N0471 (d) are shown. Positive, neutral, and negative electrostatic potentials are shown in blue, white, and red, respectively. Citrate, bis-tris propane, and cadmium ions are shown as in (b).

The C0471 domain shows strong structural homology to other GH33 sialidases, including those from *P. gingivalis*^38,39^ and *C. perfringens*^40-42^. Its conserved active site cleft (**Fig. 7a**), located at the center of the β-propeller, contains Tyr-463 and Glu-338 as the catalytic nucleophile and general base, Asp-165 as the general acid/base for water activation, and three conserved arginine residues (Arg-140, Arg-354, and Arg-423) that stabilize the carboxylate group of sialic acids, with Arg-140 forming part of the conserved RIP motif. GH33 enzymes typically contain four or five Asp-box motifs which maintain the β-propeller fold^43^. However, C0471 contains only three (**Supplementary Fig. 3**). Additionally, C0471 exhibits a pronounced electrostatic asymmetry, with a positively charged catalytic face and a negatively charged opposite face, a feature proposed to facilitate productive orientation toward negatively charged glycoconjugate substrates^44-46^. These results indicate that GH33 sialidases are structurally and catalytically conserved but with subtle divergence that underpin sialic acid recognition and catalysis.

**Figure 7.**
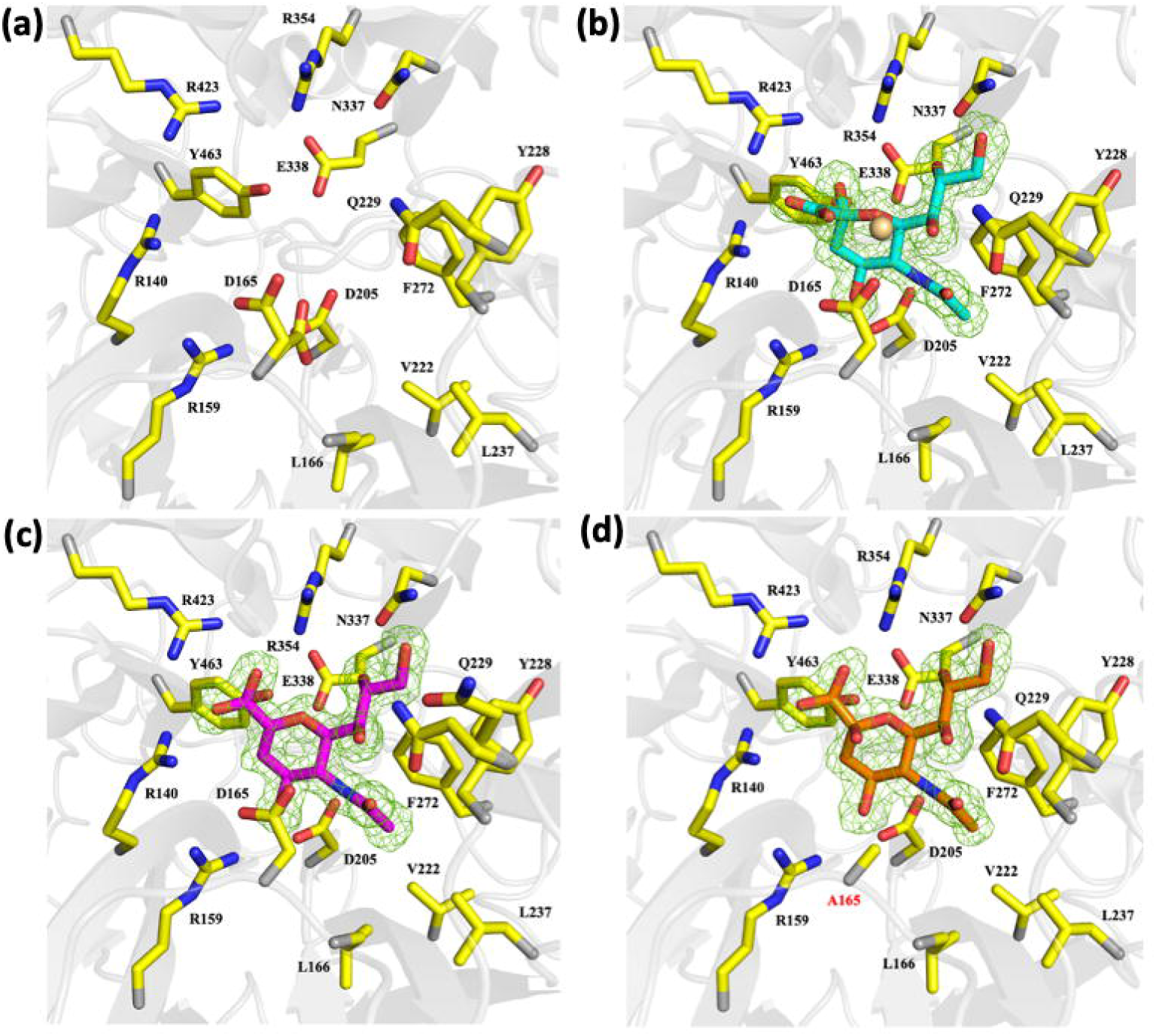
*Td*-NanH active site architecture. Cartoon representation of the sialidase active site in the (**a)** unliganded, **(b)** β**-**Neu5Ac-bound, **(c)** DANA-bound, and **(d)** 3’-SL-bound states. F_o_-F_c_ omit electron density maps, contoured at 3**σ**, for each ligand in **(b-d)** are shown as green mesh. Residues identified as belonging to the active site are shown as sticks with carbon, nitrogen, and oxygen atoms colored in yellow, blue, and red, respectively. Cadmium ions are shown as wheat spheres. In panel **(d)**, D165A mutation is labeled in red.

### Structural analyses of *Td*-NanH with sialidase ligand, inhibitor and native glycans

To define substrate interactions within the active site, we solved the crystal structures of *Td*-NanH in complex with Neu5Ac (**Fig. 7b**) and 2,3-dehydro-2-deoxy-*N*-acetylneuraminic acid (DANA, a transition-state analogue sialidase inhibitor^47^) (**Fig. 7c**) at 1.56 Å and 1.55 Å resolution, respectively. Both complexes are highly similar to the unliganded structure. Clear electron density revealed binding of β-Neu5Ac in the active site (**Fig. 7b** and **Supplementary Fig.13 & 14**), where its C4 hydroxyl forms hydrogen bonds with Arg-159, Asp-165, and Asp-205 and its N-acetyl group inserts into a hydrophobic pocket formed by Leu-166, Val-222, Tyr-228, Leu-237, and Phe-272, and is further stabilized by an ionic interaction with Asp-205. The glycerol side chain is coordinated by Gln-229 and Asn-337. Structural comparison with other two GH33 sialidase-Neu5Ac complex structures in the PDB (1E8U^48^ and 2BER^49^) indicates that the sugar ring, N-acetyl, and glycerol moieties adopt highly conserved conformations (**Fig. 8a-c**). DANA binds in an almost identical pose, with the key difference being a more planar sugar ring and carboxylate group, consistent with its role as a transition-state mimetic (**Fig. 7c**).

**Figure 8.**
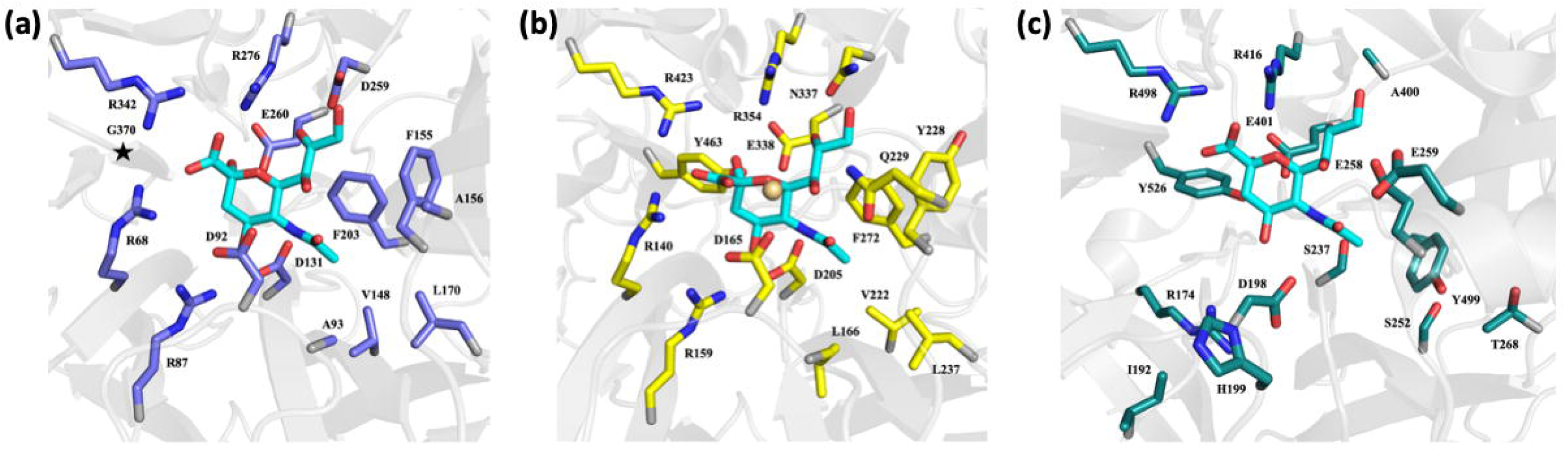
Comparison of PDB deposited sialidases bound to β-Neu5Ac. Cartoon representations of the active sites of **(a)** *Micromonospora viridifaciens* Y370G inverting sialidase mutant (PDB entry 2BER), **(b)** *Td-*NanH sialidase, and **(c)** paramyxovirus sialidase (PDB entry 1E8U) bound to β-Neu5Ac. β-Neu5Ac is shown as sticks with carbon, nitrogen, and oxygen atoms colored cyan, blue, and red, respectively. Side chains for each sialidase are shown as sticks and colored as slate (*M. viridifaciens*), yellow (*Td*-NanH), and deep teal (paramyxovirus) for carbons, while nitrogen and oxygens are colored blue and red, respectively. In **(a)** Y370G mutation is represented by a star icon.

To investigate how *Td*-NanH engages native glycans, we generated a catalytically impaired D165A mutant for co-crystallization experiments with 3’-sialyllactose (3’-SL) and 6’-sialyllactose (6’-SL). However, we were only able to grow crystals in the presence of 3’-SL and subsequently solved its structure at 1.81 Å resolution (**Fig. 7d** and **Supplementary Fig. 15**). The mutant-substrate complex superimposes closely with the unliganded-, Neu5Ac-, and DANA-bound structures (Cα RMSD 0.22– 0.26 Å), indicating that ligand binding does not induce major conformational changes. Comparison with the *Td*-NanH:DANA structure shows that active site residues are essentially identical, aside from the absence of the Asp-165 side chain (**Fig.7d**). Importantly, the terminal sialic acid of 3′-SL binds in nearly the same conformation as DANA, with only a slightly less distorted boat conformation of the sugar ring. These observations support a conserved binding mode for both free sialic acid and glycan-linked substrates and provide structural insight into the catalytic mechanism of GH33 sialidases.

In summary, our structural analyses using *Td*-NanH as a representative model demonstrate that GH33 sialidases employ a highly conserved catalytic architecture that accommodates both free and glycan-linked sialic acids in a common, transition-state-like conformation. Structural comparisons across unliganded, substrate-bound, and inhibitor-bound states reveal a rigid active-site scaffold, while subtle structural and electrostatic plasticity likely fine-tunes substrate engagement and specificity.

## DISCUSSION

Bacterial pathogens have evolved diverse strategies to evade complement-mediated killing, including proteolytic inactivation of key components such as C3, recruitment of negative regulators (for example, FH), decoration of surface structures such as lipopolysaccharide with host-derived sialic acids to dampen complement activation, and physical shielding by capsules or thick cell walls^50-52^. Here, we identify a previously unrecognized complement evasion mechanism mediated by bacterial sialidases through direct desialylation of key complement factors. To assess the generality of this strategy, we selected six sialidases from five phylogenetically and ecologically diverse bacterial pathogens. All six enzymes exhibited conserved activity toward sialylated complement proteins, resulting in enhanced resistance to complement-dependent serum killing. These findings reveal a broadly conserved and previously underappreciated mechanism of complement evasion that operates across distinct bacterial taxa and infection niches, highlighting sialidases as versatile immune-modulatory virulence factors.

Beyond complement evasion, sialidase-mediated removal of sialic acids from human serum and complement proteins may perturb immune homeostasis and exacerbate disease pathogenesis. Consistent with this notion, loss of Fc sialylation on IgG promotes pathogenic autoantibody responses^17,53^, and desialylation of FH by *S. pneumoniae* sialidase compromises its regulatory activity, leading to excessive complement activation and pathogenic effects^54^. In addition to complement components, many immune receptors critical for host defense and immune regulation, including Toll-like receptors (TLR2 and TLR4)^55,56^ and immune checkpoint molecules such as PD-1 and PD-L1^57-60^, are also modified by sialoglycans. TLR4 carries α2,6-linked Neu5Ac and PD-L1 bears α2,3-linked Neu5Ac, both of which are essential for their proper function^58,61-64^. Notably, desialylation of TLR4 by human Neu1 sialidase causes aberrant immune activation and altered responses to LPS^61,62^. Sialylation stabilizes PD-L1 and enhances its immuno-suppressive activity^64,65^. Given that complement proteins and these immune receptors share similar sialoglycan modifications, it is plausible that bacterial sialidases may also target these sialylated immune receptors and regulators, thereby modulating their function and contributing to pathogenesis. Addressing this possibility may open new avenues to define the pathogenic roles of bacterial sialidases and to exploit them as molecular tools for dissecting the largely unexplored functions of sialylation in immune regulation.

Sialylation in many complement factors remains incompletely defined^7^. Here, we show that a combined approach using sialidases, lectin blotting, and nanoLC-ESI/MS/MS provides a powerful strategy to address this gap. Applying this workflow to IgG, C1q, C4, C5, and FH, we identified multiple glycosylation sites and sialoglycans (**Fig. 1f-g, 4f-g, Supplementary Fig. 2, 11** and **Table 2-4 & Table 6-7**) covering majority of N-linked consensus sequence, despite partial sequence coverage (49-70%) obtained in each protein. Most detected N-linked glycans corresponded to previously reported structures^7,14,34,66,67^, whereas others appear to be novel, including C4 bearing HexNAc(6)Hex(3)Fuc(2) at Asn226 and C1q bearing HexNAc(4)Hex(5)Fuc(3)Neu5Ac(1) at Asn146, underscoring the complexity and heterogeneity of serum glycoproteins. We acknowledge that these findings require further validation. As sequence coverage improves through in-depth mass spectrometric analyses, this approach is likely to uncover additional glycans with relatively low abundance and further refine our understanding of complement sialylation.

Structural comparison reveals that the six tested sialidases share a conserved six-bladed β-propeller catalytic core. Despite this structural similarity, biochemical analyses demonstrate markedly different desialylation activities against complement proteins and 4-MUNANA. For example, *Td*-NanH exhibits the highest enzymatic activity against 4-MUNANA, whereas *Tf*-NanH shows the lowest. To understand the basis for these functional differences, we first examined the N-terminal regions, which vary considerably among the six sialidases and are largely uncharacterized (**Fig. 2**). Previous studies demonstrated that CMB40, an N-lectin-like domain in *Vc*-NanH, binds sialic acids and enhances its catalytic efficiency^68,69^. Like *Vc*-NanH, *Td*-NanH contains an N-terminal domain with undefined function; however, deletion of this domain had no impact on its activity against either 4-MUNANA (**Fig.5c** and **Supplementary Table 5**), suggesting that N-terminal domains do not account for the observed catalytic differences.

We therefore conducted detailed active-site comparisons using *Td*-NanH as a structural reference. Although the overall folds are highly conserved, notable differences were observed in regions coordinating the N-acetyl and glycerol moieties of Neu5Ac. In *Td*-NanH, the glycerol moiety is coordinated by Asn337; however, this residue is poorly conserved across the aligned sialidases, differing in both identity and spatial positioning (e.g., Gln493 in *Cp-*NanI and Asn425 in *Tf*-NanH; **Supplementary Fig. 17**). Similarly, the hydrophobic pocket accommodating the N-acetyl group varies substantially in residue composition across enzymes. These active-site variations likely confer differential substrate recognition and catalytic efficiency, imparting functional plasticity that fine-tunes substrate engagement and specificity across diverse sialylated targets.

While the N-terminal domain of *Td*-NanH does not contribute to its catalytic activity, we resolved citrate/bis-tris propane molecules and DANA bound in similar positions within this domain (**Supplementary Fig. 12**). Intriguingly, *T. denticola* produces a novel glycan (Td-SIA, **Supplementary Fig. 16F**) that modifies its flagellins^70^, which appears to be derived from Neu5Ac. Indeed, when the bound DANA molecule is modeled as Td-SIA in the β-anomer (**Supplementary Fig. 14**), there are no obvious clashes with *Td*-NanH, consistent with the experimentally identified β-linkage to Ser or Thr. Thus, binding of citrate/bis-tris propane and DANA raises the possibility that the N-terminal domain may enable *Td*-NanH to cleave or modify specific substrates such as Td-SIA. Additional experimentation will be required to test this hypothesis.

In summary, this study identifies complement desialylation as a conserved immune evasion strategy employed by diverse bacterial pathogens. By removing terminal sialic acids from complement proteins, GH33 sialidases impair complement activation and MAC formation, thereby promoting bacterial survival in serum. More broadly, our findings establish bacterial sialidases as multifunctional virulence factors capable of reshaping host immune responses through targeted glycan remodeling and provide a framework for the rational, structure-based development of sialidase inhibitors as anti-virulence therapeutics.

## EXPERIMENTAL PROCEDURES

### Bacterial strains, culture conditions, and oligonucleotide primers

*T. denticola* ATCC 35405 (wild type, WT) was used in this study^71^. Bacterial cells were grown in tryptone-yeast extract-gelatin-volatile fatty acids-serum (TYGVS) medium at 37°C in an anaerobic chamber in the presence of 90% nitrogen, 5% carbon dioxide, and 5% hydrogen ^72^. *T. denticola* sialidase-deficient mutants (Δ*Tde471*) were grown with erythromycin (50 µg/ml). *Escherichia coli* DH5α strain (New England Biolabs, Ipswich, MA) was used for DNA cloning; BL21 Star (DE3) (Thermo Fisher Scientific, Waltham, MA) and Lemo21(DE3) (New England Biolabs) were used for preparing recombinant proteins. *E. coli* strains were grown in lysogeny broth (LB) supplemented with appropriate concentrations of antibiotics for selective pressure as needed: ampicillin (100 µg/ml). The oligonucleotide primers for PCR used in this study are listed in **Supplementary Table 1**. These primers were synthesized in IDT (Integrated DNA Technologies, Coralville, IA).

### Electrophoresis and immunoblotting analyses

Sodium dodecyl sulfate-polyacrylamide gel electrophoresis (SDS-PAGE) and immunoblotting analyses were carried out as previously described ^28^. For immunoblotting, *T. denticola* cells were harvested in the mid-logarithmic (log) phase (∼5×10^8^ cells/ml). Equal amounts of whole-cell lysates (∼5 μg) were separated on SDS-PAGE and then transferred to polyvinylidene difluoride (PVDF) membranes. Immunoblots were probed with specific antibodies against *T. denticola* cells^28^, C3 (ab48611, Abcam, Cambridge, MA), or C9 complement protein (ab17931, Abcam). Immunoblots were developed using a horseradish peroxidase-conjugated secondary antibody with enhanced chemiluminescence (ECL) assays; signals were imaged using the Molecular Imager ChemiDoc system (Bio-Rad Laboratories, Hercules, CA), as previously described^73^.

### Lectin blot analysis

This study was performed as previously described^28^. Briefly, samples were subjected to SDS-PAGE and then transferred to PVDF membranes. Blots were first blocked in 1×Carbo-Free blocking solution (Vector Laboratories, Newark, CA) diluted in PBS-T buffer (PBS containing 0.05% Tween-20) for 1 h at room temperature. Membranes were then probed against the following lectins (Vector Laboratories and EY Laboratories), including *Sambucus nigra* (SNA, 0.2 μg/ml), Concanavalin A (ConA, 0.5 μg/mL) or *Maackia amurensis* II (MAL II, 2 μg/mL), in 1x Carbo-Free blocking buffer for 1 hour at room temperature. Blots were washed four times with PBS-T buffer and then incubated with streptavidin-horseradish peroxidase conjugate (Cytiva, Marlborough, MA). After incubation, the blots were washed four times with PBS-T buffer and developed with the ECL assay. Signals were quantified using the Molecular Imager ChemiDoc system with the Image Lab software (Bio-Rad). For lectin blot analysis of human complement proteins, the following purified proteins were used: C4b-binding protein (C4BP, 1.3□µg), Factor I (FI, 1.3□µg), Factor H (FH, 1.3□µg), C4 (2.1□µg), C5 (6.7□µg), and C1q (1.3□µg) (Complement Technologies, Tyler, TX; catalog numbers A109, A138, A137, A105, A120, and A099, respectively), as well as human immunoglobulin G (IgG, 3.8□µg) (Sigma-Aldrich, I2511).

### Evolutionary diversification of bacterial exo-sialidases

A total of 289 protein sequences belonging to the glycoside hydrolase gene family 33 (GH33) were downloaded from the UniProt and GenBank databases. Multiple sequence alignment was performed using MUSCLE (version 5.1) (http://www.ncbi.nlm.nih.gov/pubmed/15034147) with default parameters. A maximum-likelihood tree was inferred using FastTree (version 2.1.11) (http://www.ncbi.nlm.nih.gov/pubmed/20224823) with default parameters. The resulting tree was refined using the BIOTREE utility from the BpWrapper toolkit (https://bmcbioinformatics.biomedcentral.com/articles/10.1186/s12859-018-2074-9) as follows: weakly supported branches with bootstrap values <90% were removed (using the -*D* option), nodes containing closely related sequences with branch lengths <0.1 substitutions per site were collapsed and represented by a single randomly selected sequence (using the --*trim-tips* option). The tree was re-rooted at the midpoint (using the *-m* option). An alignment of the GH33 protein sequences and the associated molecular phylogeny are available through the Dryad Digital Repository.

### Preparation of recombinant bacterial sialidases

Five recombinant bacterial sialidases were prepared in this study: *T. denticola* NanH (*Td*-NanH, 25-543 aa), *C. perfringens* NanH (*Cp*-NanH, 1-382 aa), *B. fragilis* NanH2 (*Bf*-NanH2, 23-544 aa), *T. forsythia* NanH (*Tf*-NanH, 1-539 aa), and *V. cholerae* NanH (*Vc*-NanH, 94-851 aa). Nucleotide sequences encoding these sialidases were codon-optimized, synthesized (GenScript, Piscataway, NJ), and cloned into pUC19 vectors flanked by restriction sites: BamHI and HindIII for *Td*-NanH, or BamHI and SalI for the other sialidases. The codon-optimized genes were excised by digestion with BamHI/HindIII or BamHI/SalI and subcloned into the pQE-80L expression vector (Qiagen, Valencia, CA). The resulting expression constructs were transformed into BL21 Star (DE3) for *Tf-*NanH or Lemo21(DE3) for the other sialidases for protein expression and purification. A C-terminal fragment of *Td*-NanH (*Td-C*, 125-543 aa) was also prepared. The DNA sequence encoding Td-C was PCR-amplified using PrimeSTAR GXL DNA polymerase (Takara Bio USA, San Jose, CA) with primer pair P_1_/P_2_. The resulting PCR product was cloned into pJET1.2 (Thermo Fisher Scientific) and subsequently subcloned into the pQE-80L expression vector using BamHI and HindIII restriction sites. The resulting plasmid was transformed into Lemo21(DE3) for protein expression and purification. Primers used are listed in **Supplementary Table 1**.

For protein expression, 1 L of LB medium was inoculated with 25 ml of overnight *E. coli* culture and incubated at 37°C until reaching an OD□□□ of 0.4-0.8. Protein expression was induced by addition of isopropyl β-D-thiogalactopyranoside (IPTG) to a final concentration of 1 mM, and cultures were shifted to 16°C for 16-18 h. Cells were harvested by centrifugation and stored at −80°C. Recombinant proteins were purified using Ni-NTA agarose (Qiagen) under native conditions and dialyzed overnight at 4°C against phosphate-buffered saline (PBS, pH 7.4) using 3.0 kDa molecular weight cutoff Spectra/Por dialysis membranes (Spectrum Laboratories, Rancho Dominguez, CA). Protein concentrations were determined using a Bio-Rad Protein Assay Kit (Bio-Rad).

### Site-directed mutagenesis

Site-directed mutagenesis was performed using a Q5 site-directed mutagenesis kit (New England Biolabs, Ipswich, MA) or Platinum SuperFi II DNA Polymerase (Invitrogen) according to the manufacturer’s instruction. The plasmid that expresses *Td*-NanH recombinant protein was used as a template to replace either Arg140, Pro142, or Asp165 with Ala, using primers P_3_/P_4_, P_5_/P_4_, or P_6_/P_7_, respectively. Primers P_8_/P_4_ were used to replace both Arg140 and Pro142 with Ala. The resultant mutations were confirmed by DNA sequencing analysis. The primers used here are listed in **Supplementary Table 1**.

### Sialidase kinetic analysis

Kinetic study was performed on 96-well black/clear bottom plates as previously described ^28^. Briefly, 50 nM purified bacterial sialidases were incubated with increasing concentrations (15.625 - 1250 µM) of either 2′-(4-methylumbelliferyl)-α-D-*N*-acetylneuraminic acid (4-MUNANA; GoldBio, St. Louis, MO) or 4-methylumbelliferyl α-*N*-glycolylneuraminic acid (4-MUNGNA; Sussex Research, Ontario, Canada) in the appropriate reaction buffer. Reaction buffers were as follows: 100 mM sodium citrate (pH 5.0) for *Td*-NanH, *Cp*-NanH, *Cp*-NanI (N2133; MilliporeSigma, Burlington, MA), and *Bf*-NanH2; 25 mM sodium acetate, 75 mM NaCl, and 50 mM CaCl□ (pH 5.5) for *Vc*-NanH; and 35 mM sodium acetate (pH 5.5) for *Tf*-NanH. Reactions were performed at 37 °C for 1 min, and sialidase activity was immediately quantified using a Varioskan LUX multimode microplate reader (Thermo Fisher Scientific) with excitation at 365 nm and emission at 445 nm. Enzymatic activity was calculated from a 4-methylumbelliferone (4-MU; MilliporeSigma) standard curve (0-100 µM) and expressed as μmol 4-MU released per minute. Saturation curves were fitted to the Michaelis–Menten model using GraphPad Prism 10.4.1 (GraphPad Software, San Diego, CA), and kinetic parameters (K_m_ and V_max_) were determined.

### Serum killing protection assays

Normal human serum (NHS; BioIVT, Westbury, NY) or heat-inactivated serum (HIS; 56 °C for 30 min) (25 µl) was pretreated with a total of 1.7 µM purified recombinant sialidases at 37 °C for 1, 2, or 3 h in an anaerobic chamber. Sialidases were diluted in the following buffers: 50 mM sodium acetate, 150 mM NaCl, and 100 mM CaCl□ (pH 5.5) for *Vc*-NanH; or 70 mM sodium acetate (pH 5.5) for *Td*-NanH, *Cp*-NanI, *Cp*-NanH, *Bf*-NanH2, and *Tf-*NanH. Treated serum samples were then co-incubated with 25 µl mid-log-phase cultures of Δ*Tde471* (2.5 × 10□ spirochetes diluted in TYGVS medium) at 37 °C for 30 min under anaerobic conditions. Following incubation, viable spirochetes were enumerated using a Petroff–Hausser counting chamber (Hausser Scientific, Horsham, PA). Cell viability was assessed based on motility and cell surface integrity during microscopic observation (≥1 min). Survival rates were calculated as the ratio of viable cells in NHS to those in HIS, and data are presented as the mean ± standard error of the mean (SEM). Statistical significance was determined using two-way ANOVA followed by Tukey’s multiple-comparison test, with *P* < 0.01.

### Complement deposition assays

This study was performed as previously described, with minor modifications. Briefly, 50 µl of normal human serum (NHS; BioIVT) was pretreated with a total of 5 µg of either wild-type *Td*-NanH, the catalytically inactive point mutant *Td*-NanH R140A/P142A, or PBS (control) at 37 °C for 1, 2, or 3 h in an anaerobic chamber. Treated serum samples were then co-incubated with 50 µl of mid-log-phase cultures of the *T. denticola* sialidase-deficient mutant (Δ*Tde471*; 5 × 10□ spirochetes diluted in TYGVS medium) at 37 °C for 15 min under anaerobic conditions. Following incubation, samples were placed on ice for 1 min to terminate complement activation, and bacterial cells were harvested by centrifugation. Cell pellets were washed twice with ice-cold PBS and subjected to SDS–PAGE, followed by immunoblot analysis using antibodies against C3 (Abcam), C9 (Abcam), or *T. denticola*.

### Treatment of human serum and complement factors with recombinant sialidases

This assay was performed as previously described. Briefly, NHS (BioIVT) was diluted to 2% (v/v) and incubated with bacterial recombinant sialidases (ranging from 0.34 to 1 µM) at 37 °C. Samples were collected at 1 and 3 h. Reactions were performed in enzyme-specific buffers as described above. Treated samples were analyzed by SDS–PAGE followed by lectin blotting with SNA, MAL II, or ConA. Purified human complement proteins, including C1q (146.34 nM), C4 (780.48 nM), C5 (1.58 µM), FH (387.09 nM), FI (681.82 nM), C4BP (46.29 nM) (Complement Technologies), and human IgG (1.93 µM; Sigma-Aldrich), were treated with sialidases under the same conditions. Samples collected at 1 and 3 h were subjected to SDS-PAGE followed by SNA lectin blotting.

### In-gel digestion, nano LC–MS/MS analysis, and data processing

Excised gel bands were cut into ∼1 mm^3^ pieces and subjected to in-gel tryptic digestion. Gel pieces were sequentially washed at room temperature with deionized water (5 min), 50 mM ammonium bicarbonate in 50% acetonitrile (ACN; 10 min, twice), and 100% ACN (5 min), then dried in a SpeedVac concentrator (SC110, Thermo Savant). Proteins were reduced with 10 mM dithiothreitol in 100 mM ammonium bicarbonate for 1 h at 60 °C and alkylated with 55 mM iodoacetamide in 100 mM ammonium bicarbonate for 45 min at room temperature in the dark. Gel pieces were washed as above (with a single 50 mM ammonium bicarbonate/50% ACN wash), dried, and rehydrated on ice with sequencing-grade trypsin (Promega; 10 ng µl□^1^ in 50 mM ammonium bicarbonate containing 10% ACN) for 20 min. Additional ammonium bicarbonate was added to cover the gel pieces, and digestion was carried out at 37 °C for 16 h.

Digestion was terminated by adding 2% formic acid (FA) and incubating for 10 min at room temperature. Peptides were extracted twice with 50% ACN/5% FA and once with 90% ACN/5% FA, with vortexing and sonication as described, and all extracts were pooled, dried in SpeedVac, and reconstituted in 0.5% FA. Samples were filtered through 0.22 µm cellulose acetate spin filters (Corning Costar Spin-X) prior to nanoLC–ESI–MS/MS analysis. Peptide analyses were performed on an Orbitrap Fusion Tribrid mass spectrometer (Thermo Fisher Scientific) equipped with a Nanospray Flex ion source and coupled to a Dionex UltiMate 3000 RSLCnano system. Peptides (5 µl) were loaded onto a PepMap C18 viper trapping column (5 µm, 100 µm × 20 mm) at 20 µl min□^1^ and separated on a PepMap C18 analytical column (2 µm, 75 µm × 25 cm) at 35 °C using a 60-min linear gradient of 5–35% ACN in 0.1% FA at 300 nl min□^1^, followed by an 8-min ramp to 90% ACN and an 8-min hold. The column was re-equilibrated for 25 min between runs. The instrument was operated in positive ion mode with a spray voltage of 1.5 kV and a source temperature of 275 °C.

Data were acquired in data-dependent acquisition mode using a 4-s “Top Speed” method with a toggled HCD and EThcDfragmentation method. Full MS scans were acquired in the Orbitrap at a resolution of 120,000 (m/z 200) over an m/z range of 350–1600. HCD MS/MS spectra were acquired in the orbitrap analyzer for precursor ions with charge states 2-3 and intensities above 10,000 with a normalized collision energy of 30%, 3 m/z quadrupole isolation, and dynamic exclusion of 35 s (± 10 ppm). EThcD MS/MS spectra were acquired in the ion trap for precursor ions with charge states 3–7. Data acquisition was controlled using Xcalibur 4.4 software (Thermo Fisher Scientific). Raw MS and MS/MS files were searched using Byonic v3.6.8 (Protein Metrics) against the *Homo sapiens* UniProt database (26,019 entries). Searches allowed up to two missed tryptic cleavages, with carbamido-methylation of cysteine as a fixed modification, methionine oxidation, and asparagine/glutamine deamidation as variable modifications. Precursor mass tolerance was set to 10 ppm, and fragment mass tolerances were 0.05 Da for HCD and 0.6 Da for EThcD spectra. A maximum of two common and rare modifications was permitted, and N-linked glycan searches were performed against a library of 309 glycans. Identifications were filtered to a 2% false discovery rate, and all spectra corresponding to N-linked glycopeptides were manually inspected to confirm assignment confidence.

### Crystallography

Plasmids encoding wild-type *Td*-NanH and point mutant D165A were transformed into Lemo21 (DE3) *E. coli*. For large-scale expression, a 1L shaker flask containing LB media was inoculated with 25mL starter culture at 37°C and grown to an OD_600_ 0.4-0.6. The cells were induced by the addition of IPTG to a final concentration of 1mM and the temperature was lowered to 18°C. Cells were harvested 16 h post-induction via centrifugation and frozen at -80°C until further use. The cell pellet corresponding to 1L of cell growth medium was resuspended in Buffer A (50mM Tris, pH 8.0, 300mM NaCl, 10mM imidazole, 0.1% Tween-20 (v/v), 0.5mg/mL lysozyme, 60U/mL benzonase) and lysed using a microfluidizer. The cell lysate was clarified by centrifugation at 40,000*g* for 20 minutes at 4□. The resulting supernatant was incubated with 5mL of pre-equilibrated Cobalt-NTA resin (Thermo Fisher) for 30 minutes at 4°C with gentle mixing. The resin was poured into a column support and washed with 10 column volumes (CV) of Buffer B (50mM Tris, pH 8.0, 300mM NaCl, 20mM imidazole). The protein was eluted from the column using 5 CV of Buffer C (Buffer B + 300mM imidazole). Elution fractions were then pooled and concentrated to 2mL using an Ultra Centrifugal Filter with a 30kDa cutoff (Millipore). The concentrated sample was applied to a HiLoad 16/600 Superdex 200 PG size-exclusion column (Cytiva) equilibrated in 20mM Tris, pH 7.5, 150mM NaCl. The peak corresponding to TDE0471 was pooled and concentrated to 25 mg/mL using an Ultra Centrifugal Filter with a 30kDa cutoff.

Initial crystallization screening, carried out utilizing commercial screening kits and the sitting-drop vapor diffusion method, identified a lead from condition G12 (0.075M Bis-Tris Propane, 0.025M Citric Acid, pH 8.2, 0.02M Cadmium Chloride, 25% PEG400) of the Berkeley Screen (Rigaku, Bainbridge Island, WA). The initial hit appeared as liquid-liquid phase separation with strong UV signal and was optimized by standard methods, including seeding. Unliganded *Td*-NanH crystals suitable for diffraction experiments were grown at 23°C in sitting drops by combining 4μL protein solution at 7 mg/mL with 2μL reservoir solution and 0.5μL of a 1:100 dilution of seed stock and equilibrating over a 1000 μL reservoir solution of 0.025M citric acid, 0.075M Bis-Tris Propane, pH 7.4, 0.02M CdCl_2_, 25% PEG400. Crystals of *Td*-NanH for N-acetyl neuraminic acid (Neu5Ac) and 2-deoxy-2,3-dehydro-N-acetyl neuraminic acid (DANA) complexes were grown at 23°C in sitting drops by combining 4μL protein solution at 7 mg/mL with 2μL reservoir solution and equilibrating over a 1000 μL reservoir solution of 0.02M CdCl_2_, 25% PEG400, 0.01 M Tris, pH 8.2 or 0.1M HEPES, pH 7.8, respectively. To generate the complex, unliganded *Td*-NanH crystals were soaked in mother-liquor containing either 15 mM Neu5Ac for 20 seconds or 25 mM DANA for 20 minutes. *Td*-NanH complexed with 3’-sialyllactose (3’-SL) were generated via co-crystallization in 0.1M Bis-tris, pH 6.5, 0.02M CdCl_2_, 23% PEG400 with 3’-SL supplemented to the drop at 5mM. All crystals were subsequently harvested, cryoprotected in mother-liquor with 18% ethylene glycol (unliganded *Td*-NanH, *Td*-NanH:Neu5Ac, *Td*-Nan:DANA) or 20% glycerol (*Td*-NanH:3’-SL), and plunge-frozen in liquid nitrogen for diffraction analysis.

X-ray diffraction data were collected at 100K on beamlines 23-ID-B (unliganded *Td*-NanH, *Td*-NanH:Neu5Ac, and *Td*-NanH:DANA) and 23-ID-D (*Td*-NanH:3’-SL) at the Advanced Photon Source (Argonne National Laboratory) using Dectris Eiger-16M and Pilatus3 6M detectors, respectively. Data were processed with *Xia2/DIALS*^74-77^ (v3.8.0) and then scaled and merged with *Aimless* and *Pointless* ^78,79^ (v0.7.9 and v1.12.14, respectively) in the CCP4 (v8.0) suite of programs^80^. The structure of unliganded *Td*-NanH was determined by molecular replacement using *Phaser*^81^ (v2.3.8) and search models derived from a predicted structure of TDE0471 (UniProt ID: Q73QH2) generated by AlphaFold2 ^82^ and downloaded from the AlphaFold DB^83^. Two search models were generated from the original AlphaFold model, one encompassing the N-terminal domain and one comprising the sialidase domain. Each individual domain was located during the search, with the asymmetric unit containing one complete *Td*-NanH protein. The structure was initially rebuilt using *Buccaneer* ^84,85^. Successive rounds of manual model building in *Coot*^86^ and refinement in *Refmac*^87-91^ were used to build in missing pieces of the protein and ligands. In the final rounds of refinement, waters were added and translation-libration-screw refinement^92^ was applied. Structures bound with Neu5Ac, DANA, and 3’-SL were determined utilizing the same method as outlined above, except *Refmacat*^93^ was used for refinement of TDE0471:Neu5Ac, due to the presence of a cysteine sulfenic acid (Cys516). Data collection and refinement statistics for all four structures are summarized in **Supplementary Table 9**. Structure validation was performed with *MolProbity*^94^. Coordinates and structure factors have been deposited in the Protein Data Bank: unliganded *Td*-NanH (entry 10QF); *Td*-NanH:Neu5Ac (entry 10QH); *Td*-NanH:DANA (entry 10QG); and *Td*-NanH:3’-SL (entry 10GI).

### Structural alignment analysis

Structures and models were downloaded from the PDB where available, or the associated Uniprot entry for AlphaFold models. For this analysis, we used *Td*-NanH:DANA structure (this study), AlphaFold model for *Bf-*NanH2 (Uniprot entry A0A380YS77), *Tf*-NanH (PDB entry 7QY), *Cp-*NanH (PDB entry 8UB5), *Cp*-NanI (PDB entry 5TSP), *Vc*-NanH (PDB entry 1KIT). While the *B. thetaiotaomicron* NanH2 structure is available from the PDB (entry: 4BBW) we used the AlphaFold model of *B. fragilis* due to the significant structural similarity (C_⍰_ RMSD: 0.768 over 519 equivalent residues) between the two models, with a sequence identity of 74.3%. Models were then trimmed of the N-terminal CBM domains before entry into *Gesamt* (v. 1.18) in the *CCP4* suite (v. 9.0.010) of programs for structural alignment. Alignments were inspected and figures generated in PyMOL (v. 3.1.6.1). Sequence alignment was conducted with ClustalΩ in the MPI Bioinformatics and figure generated using ESPript 3.0.

## Acknowledgement

This project is supported by DE023080 to C. Li and M.G. Malkowski; DE034063 to C. Li; and USPHS award GM095459 to N.D. Clark. We thank Elizabeth Anderson in the Proteomics and Metabolomics Facility of Cornell University for technical support and NIH SIG grant 1S10 OD017992-01 to S. Zhang for the Orbitrap Fusion mass spectrometer.

## Notes

### Competing Interest Statement

The authors have declared no competing interest.

