## Supplementary Data for "Bacterial exo-α-sialidases subvert the complement system through desialylation"

Michael G. Malkowski[2*] and Chunhao Li[1*]

**Table 1. Oligonucleotide primers used in this study**

| Primers | Sequences (5'-3') | Note [a] |
|---|---|---|
| $P_1$ | GGATCCCAAGATCTGTGGCAGTCCAAG | *Td-C* recombinant protein; [F] |
| $P_2$ | AAGCTTTTATTCTTTCCAATGCAGTTTCTC | *Td-C* recombinant protein; [R] |
| $P_3$ | GGAAGATTTTTACGCCATCCCGGCGCTGGC | Site-directed mutagenesis R140A; [F] |
| $P_4$ | GGACGGTCCGTCTTGGACTGC | Site-directed mutagenesis R140A, P142A, and R140AP142A; [R] |
| $P_5$ | GGAAGATTTTTACCGCATCGCGGCGCTGGCG | Site-directed mutagenesis P142A; [F] |
| $P_6$ | CGTTACAAGAATAACAGCGCGCTGGGCAAC AACCATCGTATCG | Site-directed mutagenesis D165A; [F] |
| $P_7$ | GGTTGTTGCCCAGCGCGCTGTTATTCTTGT AACGCAGGTCAG | Site-directed mutagenesis D165A; [R] |
| $P_8$ | GGAAGATTTTTACGCCATCGCGGCGCTGGC | Site-directed mutagenesis R140A P142A; [F] |

[a] Underlined sequences are engineered restriction cut sites for DNA cloning; [F] forward; [R] reverse.

**Table 2. Glycans detected in human C4 protein by nano LC-ESI-MS/MS**

| C4 | | |
|---|---|---|
| **Position** | **Peptide Sequences** | **Identified Glycans** |
| Asn226 | R.FSDGLESN[+1216.42286]SSTQFEVK.K | HexNAc(2)Hex(5) |
| Asn226 | R.FSDGLESN[+1540.52851]SSTQFEVK.K | HexNAc(2)Hex(7) |
| Asn226 | R.FSDGLESN[+1702.58133]SSTQFEVK.K | HexNAc(2)Hex(8) |
| Asn226 | R.FSDGLESN[+1864.63416]SSTQFEVK.K | HexNAc(2)Hex(9) |
| Asn226 | R.FSDGLESN[+1864.63416]SSTQFEVKKYVLPNFEVK.I | HexNAc(2)Hex(9) |
| Asn226 | R.FSDGLESN[+1996.75052]SSTQFEVK.K | HexNAc(6)Hex(3)Fuc(2) |
| Asn226 | R.FSDGLESN[+2026.68698]SSTQFEVK.K | HexNAc(2)Hex(10) |
| Asn1328 | R.GLN[+2204.77244]VTLSSTGR.N | HexNAc(4)Hex(5)NeuAc(2) |
| Asn1328 | R.GLN[+2204.77244]VTLSSTGRNGFKSHALQLNNR.Q | HexNAc(4)Hex(5)NeuAc(2) |
| Asn1328 | R.GLN[+2204.77244]VTLSSTGRN[+0.98402]GFK.S | HexNAc(4)Hex(5)NeuAc(2) |

| | | |
|---|---|---|
| Asn1328 | R.GLN[+2205.79284]VTLSSTGRN[+0.98402]GFK.S | HexNAc(4)Hex(5)Fuc(2)NeuAc(1) |
| Asn1328 | R.GLN[+2350.83035]VTLSSTGR.N | HexNAc(4)Hex(5)Fuc(1)NeuAc(2) |
| Asn1391 | R.TYNVLDM[+15.99492]KN[+1913.67702]TTC[+57.02146]QDLQIEVTVK.G | HexNAc(4)Hex(5)NeuAc(1) |
| Asn1391 | R.TYNVLDM[+15.99492]KN[+2204.77244]TTC[+57.02146]QDLQIEVTVK.G | HexNAc(4)Hex(5)NeuAc(2) |
| Asn1391 | R.TYNVLDM[+15.99492]KN[+2205.79284]TTC[+57.02146]QDLQIEVTVK.G | HexNAc(4)Hex(5)Fuc(2)NeuAc(1) |
| Asn1391 | R.TYNVLDM[+15.99492]KN[+2221.78776]TTC[+57.02146]QDLQIEVTVK.G | HexNAc(4)Hex(6)Fuc(1)NeuAc(1) |
| Asn1391 | R.TYNVLDM[+15.99492]KN[+2245.79899]TTC[+57.02146]QDLQIEVTVK.G | HexNAc(5)Hex(4)NeuAc(2) |

### Table 3. Glycans detected in human C1q subunit A by nano LC-ESI-MS/MS

| C1q, Subunit A | | |
|---|---|---|
| **Position** | **Peptide Sequences** | **Identified Glycans** |
| Asn146 | R.RNPPM[+15.99492]GGNVVIFDTVITNQEEPYQN[+2352.84600]HSGR.F | HexNAc(6)Hex(7) |
| Asn146 | R.RNPPM[+15.99492]GGNVVIFDTVITNQEEPYQN[+2351.85075]HSGR.F | HexNAc(4)Hex(5)Fuc(3)NeuAc(1) |
| Asn146 | R.RNPPM[+15.99492]GGNVVIFDTVITNQEEPYQN[+2059.73493]HSGR.F | HexNAc(4)Hex(5)Fuc(1)NeuAc(1) |
| Asn146 | R.NPPM[+15.99492]GGNVVIFDTVITNQEEPYQN[+2350.83035]HSGR.F | HexNAc(4)Hex(5)Fuc(1)NeuAc(2) |
| Asn146 | R.NPPM[+15.99492]GGNVVIFDTVITNQEEPYQN[+2076.75025]HSGR.F | HexNAc(4)Hex(6)Fuc(2) |
| Asn146 | R.NPPM[+15.99492]GGNVVIFDTVITNQEEPYQN[+2059.73493]HSGR.F | HexNAc(4)Hex(5)Fuc(1)NeuAc(1) |

### Table 4. Glycans detected in human IgG by nano LC-ESI-MS/MS

| IgG | | |
|---|---|---|
| **Position** | **Peptide Sequences** | **Identified Glycans** |
| Asn297 | K.TKPREEQYN[+1298.47596]STYR.V | HexNAc(4)Hex(3) |
| Asn297 | K.TKPREEQYN[+1444.53387]STYR.V | HexNAc(4)Hex(3)Fuc(1) |
| Asn297 | K.TKPREEQYN[+1460.52878]STYR.V | HexNAc(4)Hex(4) |
| Asn297 | K.TKPREEQYN[+1501.55533]STYR.V | HexNAc(5)Hex(3) |
| Asn297 | K.TKPREEQYN[+1590.59178]STYR.V | HexNAc(4)Hex(3)Fuc(2) |
| Asn297 | K.TKPREEQYN[+1606.58669]STYR.V | HexNAc(4)Hex(4)Fuc(1) |
| Asn297 | K.TKPREEQYN[+1622.58161]STYR.V | HexNAc(4)Hex(5) |
| Asn297 | K.TKPREEQYN[+1647.61324]STYR.V | HexNAc(5)Hex(3)Fuc(1) |
| Asn297 | K.TKPREEQYN[+1663.60816]STYR.V | HexNAc(5)Hex(4) |
| Asn297 | K.TKPREEQYN[+1751.62420]STYR.V | HexNAc(4)Hex(4)NeuAc(1) |
| Asn297 | K.TKPREEQYN[+1768.63952]STYR.V | HexNAc(4)Hex(5)Fuc(1) |
| Asn297 | K.TKPREEQYN[+1809.66607]STYR.V | HexNAc(5)Hex(4)Fuc(1) |
| Asn297 | K.TKPREEQYN[+1825.66098]STYR.V | HexNAc(5)Hex(5) |
| Asn297 | K.TKPREEQYN[+1897.68211]STYR.V | HexNAc(4)Hex(4)Fuc(1)NeuAc(1) |
| Asn297 | K.TKPREEQYN[+1913.67702]STYR.V | HexNAc(4)Hex(5)NeuAc(1) |
| Asn297 | K.TKPREEQYN[+1971.71889]STYR.V | HexNAc(5)Hex(5)Fuc(1) |
| Asn297 | K.TKPREEQYN[+2059.73493]STYR.V | HexNAc(4)Hex(5)Fuc(1)NeuAc(1) |
| Asn297 | R.EEQYN[+1444.53387]STYR.V | HexNAc(4)Hex(3)Fuc(1) |
| Asn297 | R.EEQYN[+1606.58669]STYR.V | HexNAc(4)Hex(4)Fuc(1) |
| Asn297 | R.EEQYN[+1622.58161]STYR.V | HexNAc(4)Hex(5) |
| Asn297 | R.EEQYN[+1663.60816]STYR.V | HexNAc(5)Hex(4) |
| Asn297 | R.EEQYN[+1768.63952]STYR.V | HexNAc(4)Hex(5)Fuc(1) |
| Asn297 | R.EEQYN[+1809.66607]STYR.V | HexNAc(5)Hex(4)Fuc(1) |

**Table 5. Kinetic analysis of bacterial exo-*α*-sialidases**

| Protein | Substrate | Vmax (µM/min ± SEM) | Km (µM ± SEM) |
|---|---|---|---|
| *Td*-NanH | 4-MUNANA | 995.2 ± 25.06 | 840.9 ± 41.70 |
| *Td*-NanH | 4-MUNAGc | 549.8 ± 36.7 | 1,370 ± 151.8 |
| *Td-C* | 4-MUNANA | 1,046 ± 32.19 | 867.9 ± 52.01 |
| *Td*-NanH$^{R140A}$ | 4-MUNANA | 234.6 ± 11.30 | 1320 ± 106.8 |
| *Td*-NanH$^{P142A}$ | 4-MUNANA | 373.7 ± 5.78 | 278.3 ± 12.48 |
| *Td*-NanH$^{R140AP142A}$ | 4-MUNANA | ND | ND |
| *Bf*-NanH2 | 4-MUNANA | 392.3 ± 8.76 | 268.9 ± 17.6 |
| *Cp*-NanI | 4-MUNANA | 720 ± 16.5 | 851.2 ± 38.24 |
| *Cp*-NanH | 4-MUNANA | 898.6 ± 30.84 | 762.6 ± 53.41 |
| *Tf*-NanH | 4-MUNANA | 96.89 ± 2.61 | 171.9 ± 15.44 |
| *Vc*-NanH | 4-MUNANA | 939.7 ± 27.75 | 606.0 ± 39.74 |

**Table 6. Glycans detected in human C5 by nano LC-ESI-MS/MS**

| Untreated C5 | | | *Td*-NanH Treated C5 | | |
|---|---|---|---|---|---|
| Position | Peptide Sequences | Identified Glycans | Position | Peptide Sequences | Identified Glycans |
| Asn741 | R.AN[+2204.77244]ISHK.D | HexNAc(4)Hex(5)NeuAc(2) | Asn741 | R.AN[+1622.58161]ISHKDM [+15.99492]QLGR.L | HexNAc(4)Hex(5) |
| Asn741 | R.AN[+2204.77244]ISHKDM [+15.99492]QLGR.L | HexNAc(4)Hex(5)NeuAc(2) | ND | ND | ND |
| Asn1630 | K.YN[+1913.67702]FSFR.Y | HexNAc(4)Hex(5)NeuAc(1) | Asn1630 | K.YN[+1622.58161]FSFR.Y | HexNAc(4)Hex(5) |
| Asn1630 | K.YN[+2204.77244]FSFR.Y | HexNAc(4)Hex(5)NeuAc(2) | Asn1630 | K.YN[+1987.71380]FSFR.Y | HexNAc(5)Hex(6) |
| Asn1630 | K.YN[+2350.83035]FSFR.Y | HexNAc(4)Hex(5)Fuc(1)NeuAc(2) | Asn1630 | K.YN[+1768.63952]FSFR.Y | HexNAc(4)Hex(5)Fuc(1) |

**Table 7. Glycans detected in Factor H protein by nano LC-ESI-MS/MS**

| Untreated Factor H | | | *Td*-NanH-Treated Factor H | | |
|---|---|---|---|---|---|
| Position | Peptide Sequences | Identified Glycans | Position | Peptide Sequences | Identified Glycans |
| | | | Asn217 | K.SPDVIN[+1622.58161]GSPIS QK.I | HexNAc(4)Hex(5) |
| Asn882 | K.IPC[+57.02146]SQPPQIEHGT IN[+1913.67702]SSR.S | HexNAc(4)Hex(5)NeuAc(1) | Asn882 | K.IPC[+57.02146]SQPPQIEHGT IN[+1622.58161]SSR.S | HexNAc(4)Hex(5) |
| Asn882 | K.IPC[+57.02146]SQPPQIEHGT IN[+2204.77244]SSR.S | HexNAc(4)Hex(5)NeuAc(2) | Asn882 | K.IPC[+57.02146]SQPPQIEHGT IN[+1913.67702]SSR.S | HexNAc(4)Hex(5)NeuAc(1) |
| | | | Asn882 | K.IPC[+57.02146]SQPPQIEHGT IN[+1987.71380]SSR.S | HexNAc(5)Hex(6) |
| Asn911 | R.ISEEN[+1913.67702]ETTC[+5 7.02146]YM[+15.99492]GK.W | HexNAc(4)Hex(5)NeuAc(1) | Asn911 | R.ISEEN[+1622.58161]ETTC[+5 7.02146]YMGK.W | HexNAc(4)Hex(5) |
| Asn911 | R.ISEEN[+2204.77244]ETTC[+5 7.02146]YMGK.W | HexNAc(4)Hex(5)NeuAc(2) | Asn911 | R.ISEEN[+1622.58161]ETTC[+5 7.02146]YM[+15.99492]GK.W | HexNAc(4)Hex(5) |

| | | | | | |
|---|---|---|---|---|---|
| Asn911 | R.ISEEN[+2204.77244]ETTC[+57.02146]YM[+15.99492]GK.W | HexNAc(4)Hex(5)NeuAc(2) | Asn911 | R.ISEEN[+1768.63952]ETTC[+57.02146]YM[+15.99492]GK.W | HexNAc(4)Hex(5)Fuc(1) |
| Asn911 | R.ISEEN[+2205.79284]ETTC[+57.02146]YM[+15.99492]GK.W | HexNAc(4)Hex(5)Fuc(2)NeuAc(1) | Asn911 | R.ISEEN[+1987.71380]ETTC[+57.02146]YMGK.W | HexNAc(5)Hex(6) |
| | | | Asn911 | R.ISEEN[+2133.77171]ETTC[+57.02146]YM[+15.99492]GK.W | HexNAc(5)Hex(6)Fuc(1) |
| Asn1029 | K.MDGASN[+1913.67702]VTC[+57.02146]INSR.W | HexNAc(4)Hex(5)NeuAc(1) | Asn1029 | K.MDGASN[+1622.58161]VTC[+57.02146]INSR.W | HexNAc(4)Hex(5) |
| Asn1029 | K.MDGASN[+2059.73493]VTC[+57.02146]INSR.W | HexNAc(4)Hex(5)Fuc(1)NeuAc(1) | Asn1029 | K.M[+15.99492]DGASN[+1460.52878]VTC[+57.02146]INSR.W | HexNAc(4)Hex(4) |
| Asn1029 | K.M[+15.99492]DGASN[+1913.67702]VTC[+57.02146]INSR.W | HexNAc(4)Hex(5)NeuAc(1) | Asn1029 | K.M[+15.99492]DGASN[+1622.58161]VTC[+57.02146]INSR.W | HexNAc(4)Hex(5) |
| Asn1029 | K.M[+15.99492]DGASN[+2204.77244]VTC[+57.02146]INSR.W | HexNAc(4)Hex(5)NeuAc(2) | Asn1029 | K.M[+15.99492]DGASN[+1768.63952]VTC[+57.02146]INSR.W | HexNAc(4)Hex(5)Fuc(1) |
| | | | Asn1029 | K.M[+15.99492]DGASN[+1913.67702]VTC[+57.02146]INSR.W | HexNAc(4)Hex(5)NeuAc(1) |

### Table 8. Sequence identity among six GH33 sialidases

| | ***Vc*-NanH** | ***Td*-NanH** | ***Cp*-NanI** | ***Cp*-NanH** | ***Bf*-NanH** | ***Tf*-NanH** |
|---|---|---|---|---|---|---|
| ***Vc*-NanH** | 100 | 17.23 | 22.33 | 23.10 | 18.79 | 18.95 |
| ***Td*-NanH** | 17.23 | 100 | 25.77 | 20.64 | 22.25 | 21.89 |
| ***Cp*-NanI** | 22.33 | 25.77 | 100 | 28.33 | 27.71 | 27.29 |
| ***Cp*-NanH** | 23.10 | 20.64 | 28.33 | 100 | 26.04 | 28.89 |
| ***Bf*-NanH** | 18.79 | 22.25 | 27.71 | 26.04 | 100 | 64.94 |
| ***Tf*-NanH** | 18.95 | 21.89 | 27.29 | 28.89 | 64.94 | 100 |

### Table 9. Crystallographic statistics for *Td*-NanH datasets.

| Dataset | Apo3 | Neu5Ac | DANA | 3SL |
|---|---|---|---|---|
| Space group | C2 | C2 | C2 | C2 |
| Wavelength (Å) | 1.033 | 1.033 | 1.033 | 1.033 |
| Unit cell length (Å) | | | | |
| a | 129.79 | 129.73 | 129.81 | 130.45 |
| b | 65.48 | 65.37 | 65.55 | 66.08 |
| c | 92.04 | 92.02 | 91.68 | 93.15 |
| β (°) | 106.78 | 106.19 | 106.15 | 106.62 |
| Resolution (Å) | 45.71 – 1.63 | 57.95 – 1.56 | 59.31 – 1.55 | 46.17 – 1.81 |
| Highest Resolution Shell (Å) | 1.67 – 1.63 | 1.60 – 1.56 | 1.59 – 1.55 | 1.86 – 1.81 |
| Total observations [a] | 243085 (10837) | 474795 (20614) | 486732 (23092) | 201571 (11706) |
| Total unique | 88676 (4083) | 102901 (4741) | 104014 (4956) | 69062 (4050) |

| | | | | |
|---|---|---|---|---|
| Multiplicity | 2.7 (2.7) | 4.6 (4.4) | 4.7 (4.7) | 2.9 (2.9) |
| Completeness (%) | 95.8 (88.5) | 97.3 (90.2) | 97.1 (94.6) | 99.7 (99.5) |
| Mean I / σ(I) | 5.7 (1.0) | 11.2 (3.0) | 11.8 (4.5) | 4.8 (1.2) |
| $R_{merge}$ (%) [b] | 10.9 (103.3) | 7.5 (42.2) | 7.7 (27.4) | 17.4 (110.8) |
| $CC^{1/2}$ [c] | 0.985 (0.571) | 0.997 (0.912) | 0.994 (0.943) | 0.979 (0.429) |
| CC* [c] | 0.996 (0.853) | 0.999 (0.977) | 0.998 (0.985) | 0.995 (0.775) |
| Wilson B-factor ($Å^2$) | 12.73 | 12.28 | 9.5 | 11.5 |
| Number of non-hydrogen atoms | 4746 | 4784 | 4914 | 4640 |
| $R_{work}$ | 0.164 (0.287) | 0.142 (0.209) | 0.138 (0.170) | 0.165 (0.291) |
| $R_{free}$ [d] | 0.187 (0.311) | 0.158 (0.229) | 0.153 (0.202) | 0.190 (0.308) |
| Average B-factor, protein ($Å^2$) | 18.46 | 20.12 | 16.14 | 18.24 |
| Average B-factor, solvent ($Å^2$) | 29.94 | 31.04 | 26.77 | 26.85 |
| Coordinate error (Å) | 0.060 | 0.037 | 0.032 | 0.080 |
| RMSD bonds lengths (Å) | 0.013 | 0.013 | 0.013 | 0.014 |
| RMSD bond angles (°) | 1.601 | 1.81 | 1.656 | 1.863 |
| Ramachandran plot | | | | |
| Favored (%) | 97.39 | 96.36 | 96.79 | 97.77 |
| Allowed (%) | 2.41 | 3.64 | 3.21 | 2.03 |
| Disallowed (%) | 0.20 | 0.00 | 0.00 | 0.00 |
| Clash score [e] | 2.04 | 2.15 | 1.18 | 1.86 |

[a] Values in parentheses represent the values in the highest resolution shell.

[b] $R_{merge}$ as defined in [1].

[c] $CC^{1/2}$ and CC* as defined in [2].

[d] Respectively, 4601, 5155, 5139, 3493 of the reflections (5.2%, 5.3%, 5.3%, 5.3%) were utilized in the test set.

[e] Clash score as calculated in *MolProbity* [3].

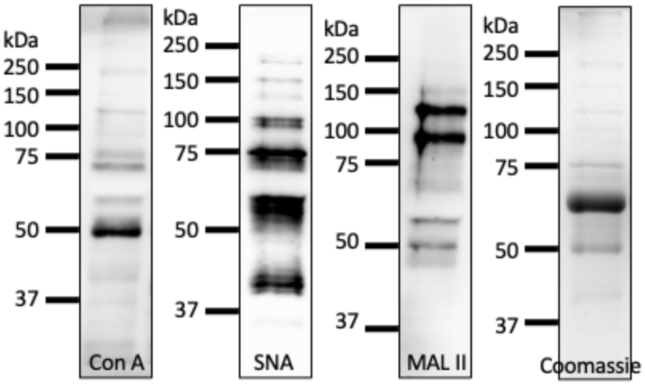


**Figure S1. Human serum proteins are glycosylated and sialylated.** 2% normal human serum (NHS) was subjected to SDS-PAGE, followed by Coomassie staining and lectin blots using Concanavalin A (ConA), *Sambucus nigra* agglutinin (SNA) and *Maackia amurensis* II (MAL II).

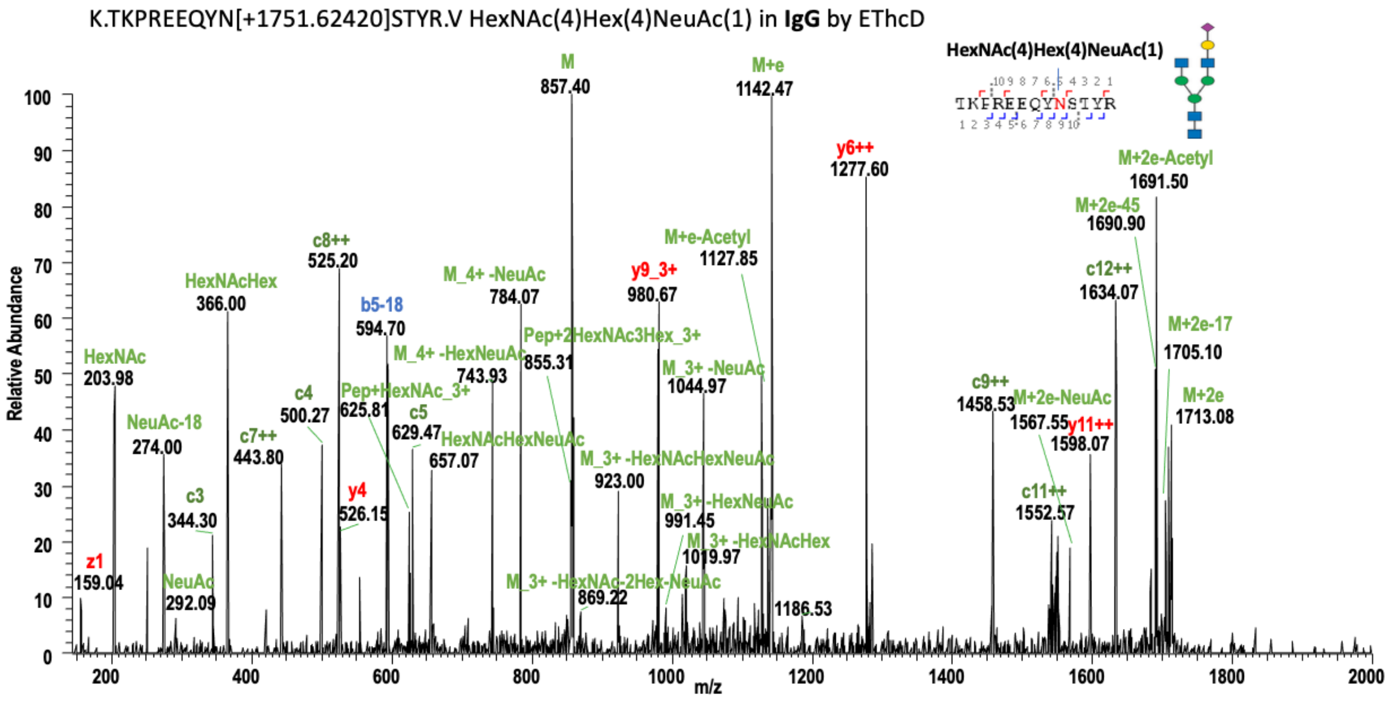


**Figure S2. A representative nano LC-ESI-MS/MS spectrum of human IgG.**

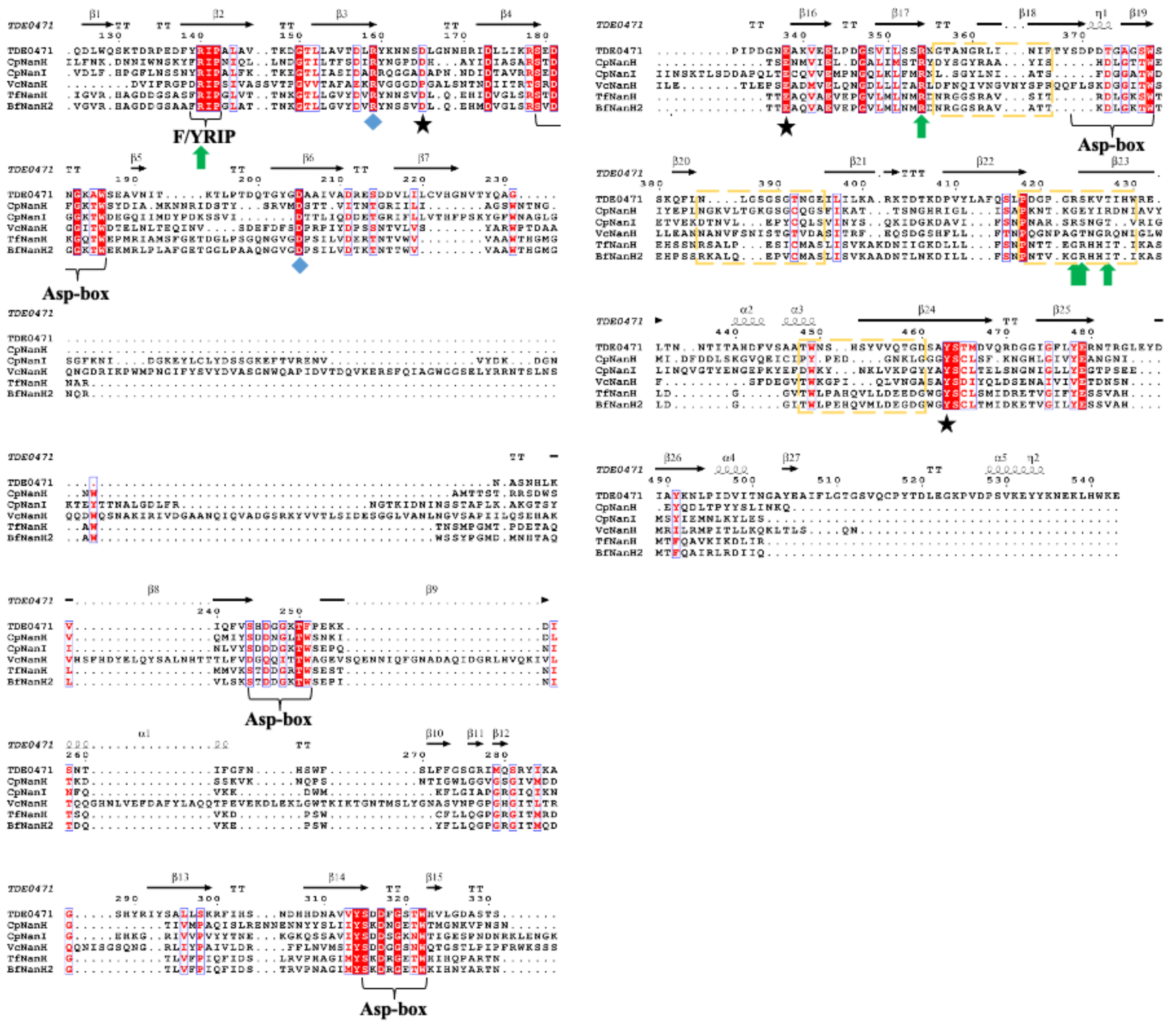

**Figure S3. Multiple sequence alignment of six sialidases.** Shown is the multiple sequence alignment (MSA) between *T. denticola* TDE0471 (*Td*-NanH), *C. perfringens* NanH and NanI (*Cp*-NanH and *Cp*-NanI, respectively), *V. cholera* NanH (*Vc*-NanH), *T. forsythia* NanH (*Tf*-NanH), and *B. fragilis* NanH2 (*Bf*-NanH2). Secondary structure corresponding to *Td*-NanH is depicted above the MSA, with beta-strands, alpha helices, $3_{10}$-helices, strict beta-turns, and strict alpha-turns depicted as arrows (β), squiggles (α), squiggles (η), "TT", and "TTT", respectively, and numbered. A red box with white character signifies strict identity, a red character signifying similarity, and "." Indicating a gap. The conserved "Y/FRIP" sequence and "Asp-box" sequences are labeled. Green arrows signify the "Tri-Arg" motif residues, catalytic residues are depicted as black stars, purple diamonds depict other conserved residues highlighted in the text. Loops previously implicated in substrate selectivity by previous studies are denoted by blue dashed boxes. The catalytic aspartic acid is shifted one residue to the left in *Vc*-NanH and the last "Tri-Arg" motif arginine is shifted to the right by one residue in *Tf*-NanH and *Bf*-NanH and four residues to the right in *Cp*-NanH and *Vc*-NanH.

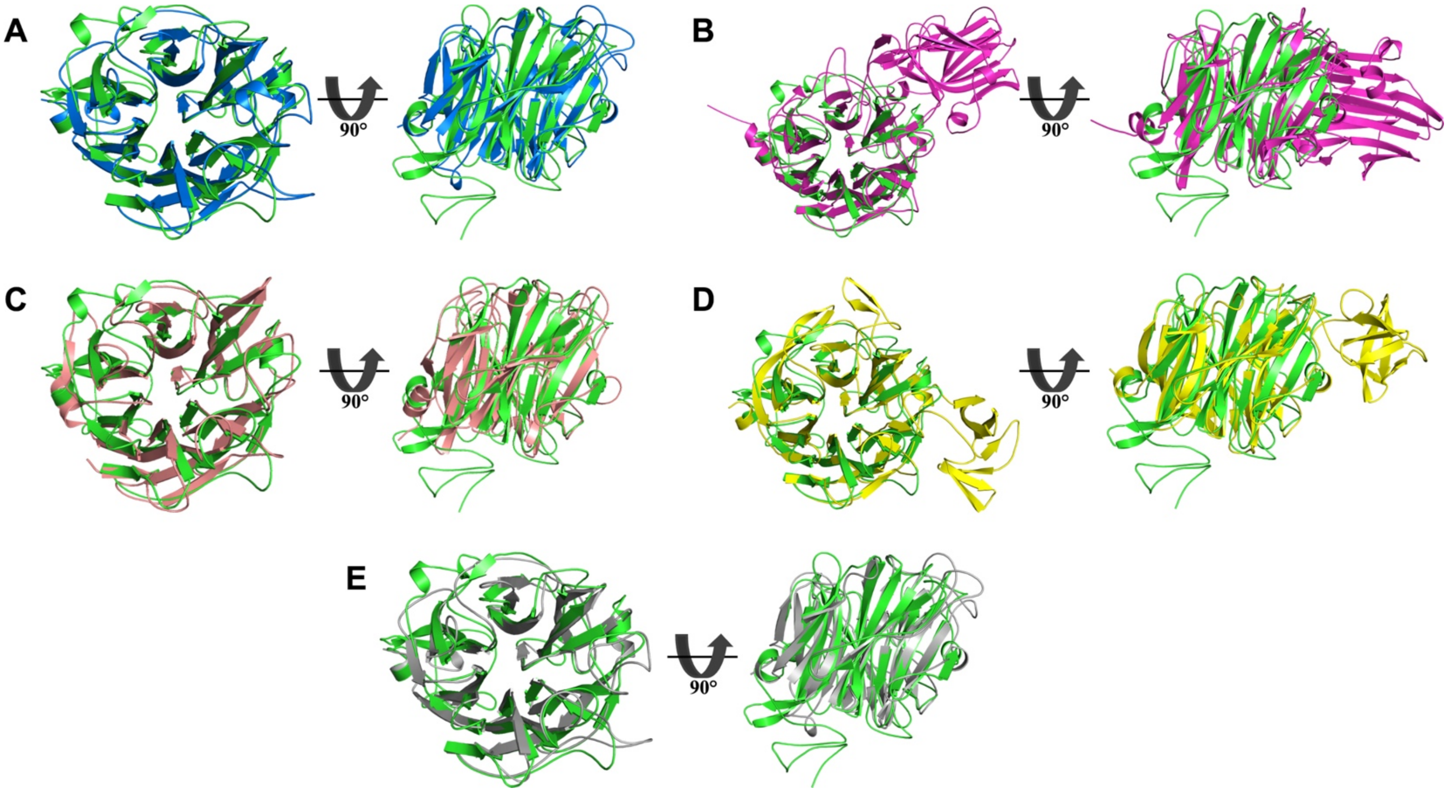


**Figure S4. Global structural alignment of six sialidases.** (A-E) Depicted is the structural overlay, shown as cartoon, between TDE0471 (green) and (A) *T. forsythia* NanH (marine; PDB entry: 7QYP), (B) *V. cholera* NanH (magenta; PDB entry: 1KIT), (C) *C. perfringens* NanH (salmon; 8UB5), (D) *C. perfringens* NanI (yellow; PDB entry: 5TSP), (E) *B. fragilis* NanH2 (grey; AlphaFold). In the top view (left panel) the active site is facing out of the page and in the side view (right panel) the active site is facing the top of the page.

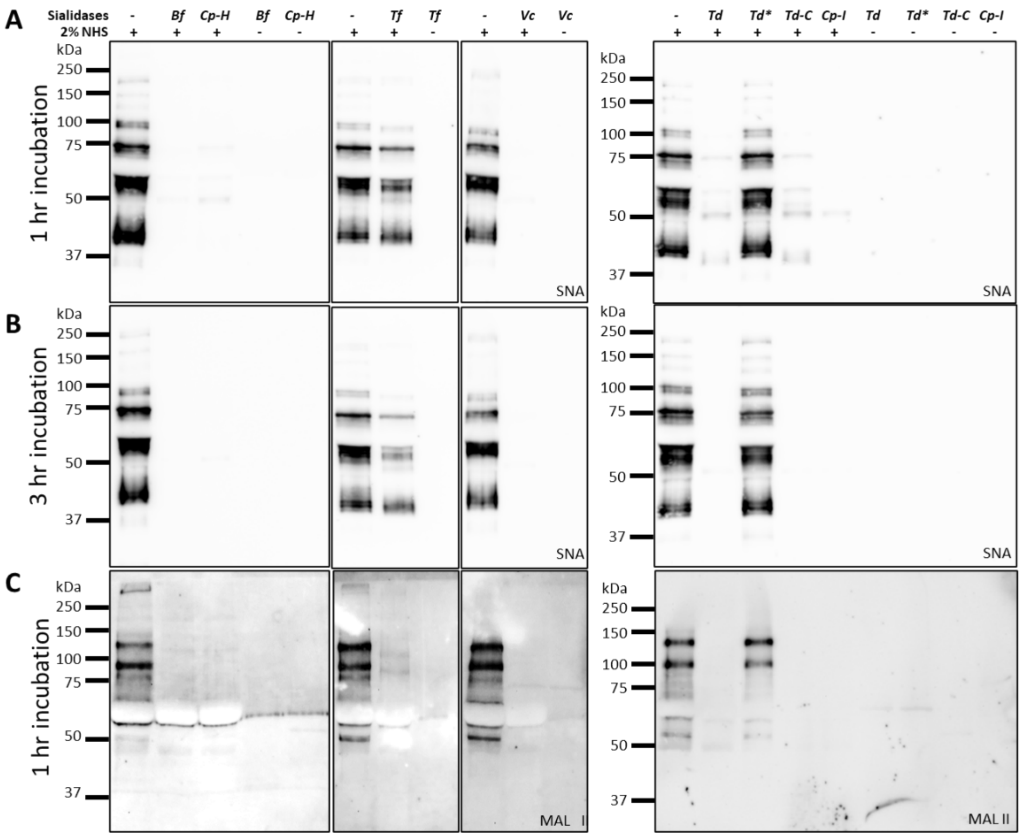

**Figure S5. GH33 sialidases remove human serum sialic acids.** Normal human serum (NHS) was incubated with GH33 sialidases at 37°C for 1 or 3 hours, followed by SDS-PAGE and lectin blot analysis using SNA (A, B) and MAL II (C) lectins. Abbreviations: *T. denticola* NanH (*Td*-NanH; *Td*), *T. denticola* NanH R140A P142A (*Td*-NanH*; *Td**), *T. denticola* NanH C-terminal (*Td-C*), *C. perfringens* NanI (*Cp*-NanI; *Cp-I*), *C. perfringens* NanH (*Cp*-NanH; *Cp-H*), *B. fragilis* NanH2 (*Bf*–NanH2; *Bf*), *T. forsythia* NanH (*Tf*-NanH; *Tf*), and *V. cholera* NanH (*Vc*-NanH; *Vc*)

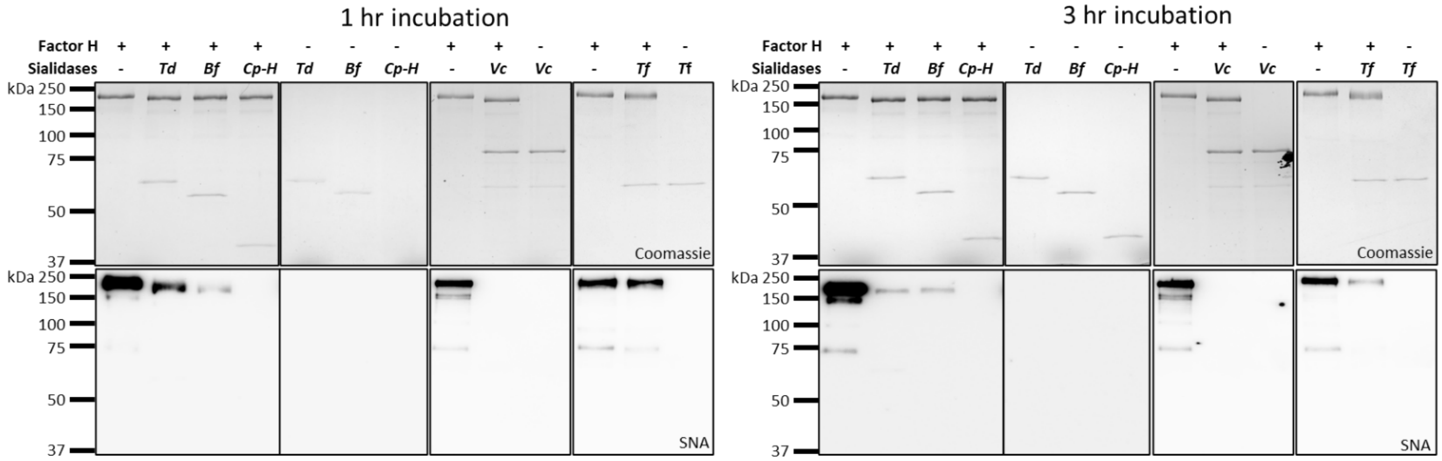


**Figure S6. GH33 sialidases desialylate complement regulatory protein Factor H (FH).** For this experiment, FH protein (387.09 nM) was incubated with six different GH33 sialidases at 37°C for 1 or 3 hours, followed by SDS-PAGE and Coomassie staining (top panel) and lectin blot analysis using SNA lectin (lower panel).

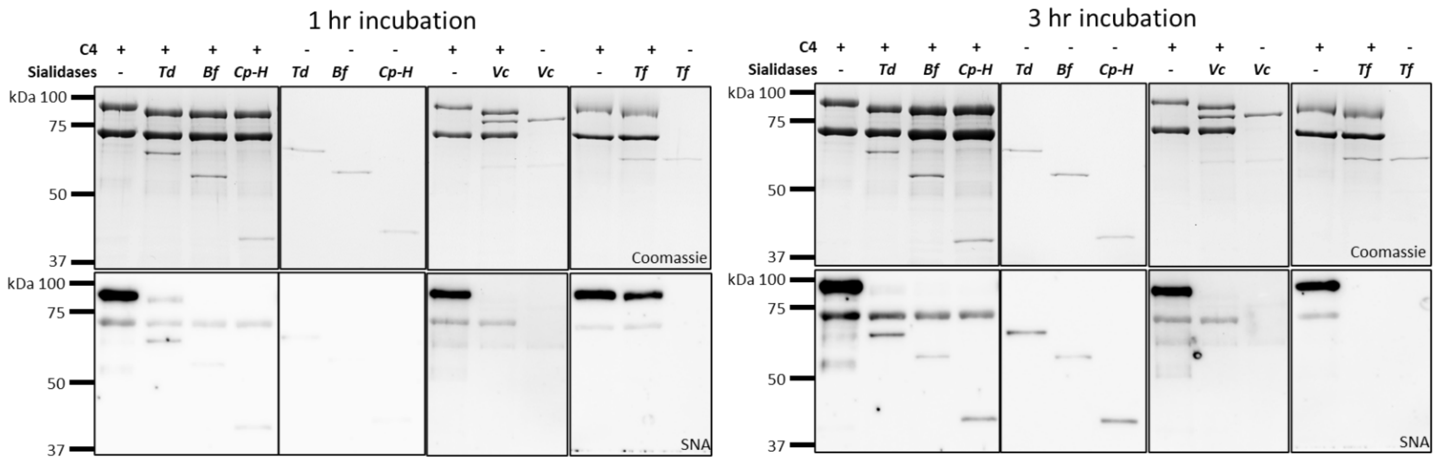


**Figure S7. GH33 sialidases desialylate complement C4 protein.** For this study, C4 protein (780.48 nM) was incubated with six different GH33 sialidases at 37°C for 1 or 3 hours, followed by SDS-PAGE and Coomassie staining (top panel) and lectin blot analysis using SNA lectin (lower panel).

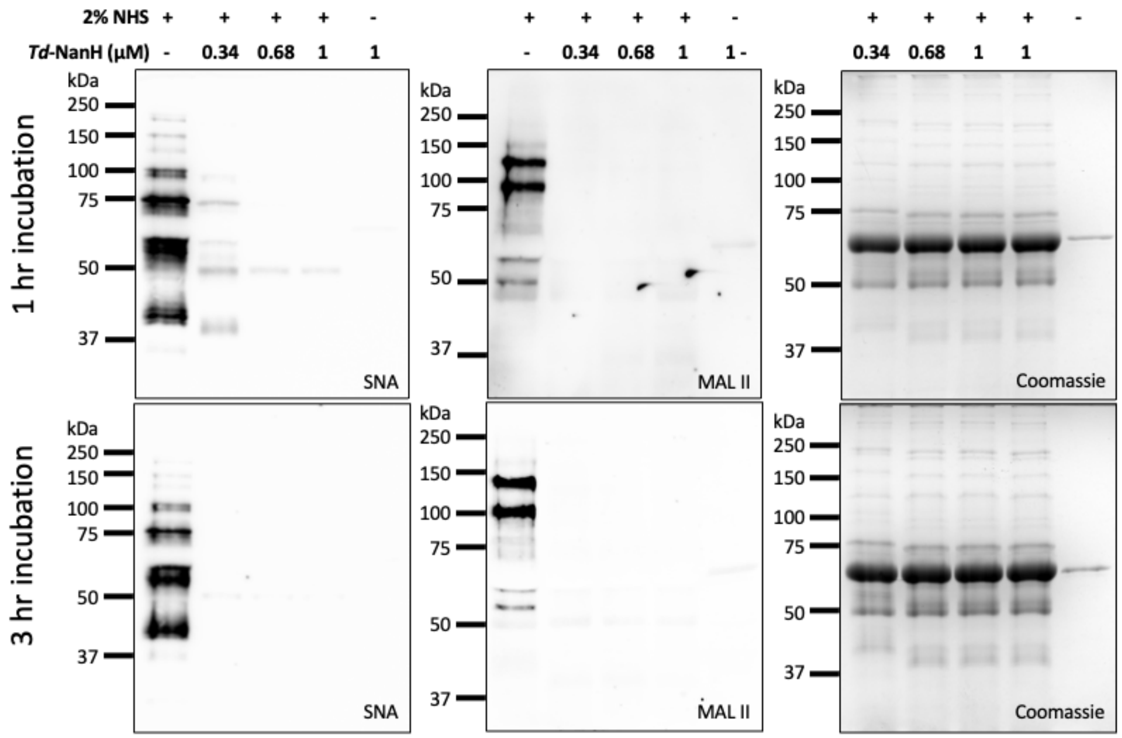


**Figure S8. *Td*-NanH desialylates human serum proteins.** Normal human serum (NHS) was treated with varying amount of *Td*-NanH as labeled at 37°C for 1 or 3 hours. Samples were analyzed by SDS-PAGE, followed by Coomassie staining and lectin blot analysis using SNA and MAL II lectins.

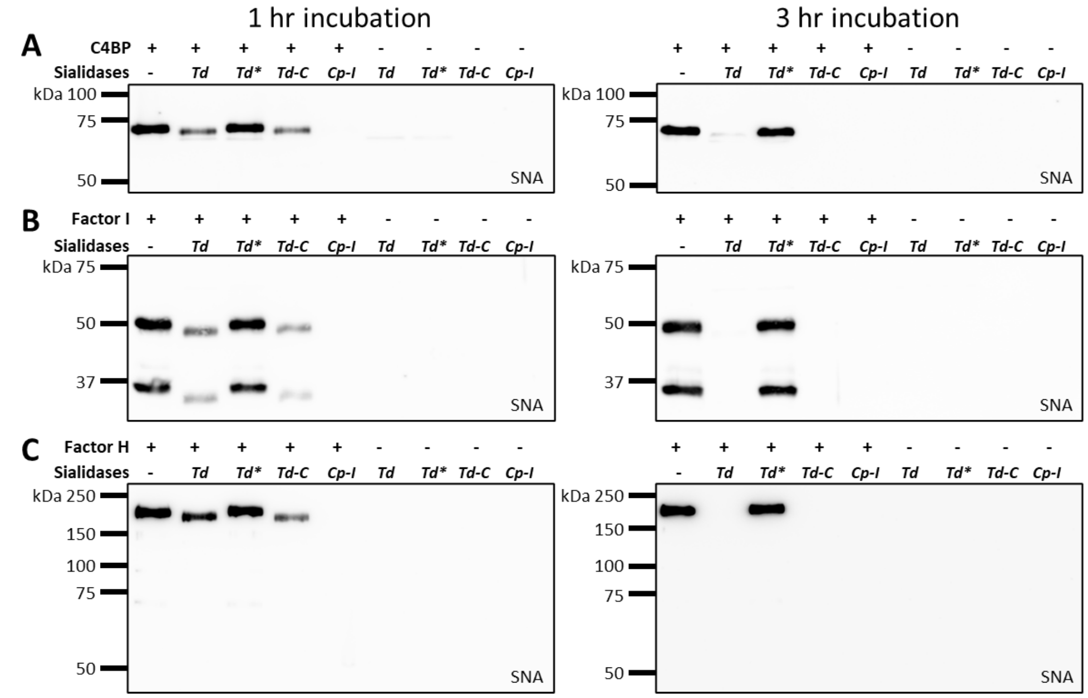


**Figure S9. *Td*-NanH desialylates complement proteins C4BP (A), Factor I (B), and Factor H (C).** For this study, three complement proteins were incubated with sialidases at 37°C for 1 or 3 hours and then subjected to SDS-PAGE followed by lectin blots using SNA lectin. *Td,* the wild type *T. denticola* NanH*; Td**, an catalytic inactive mutant of *Td*-NanH (R140AP142A); *Td-C*; the C-terminal fragment of *Td*-NanH; and *Cp-I*, *C. perfringens* NanI.

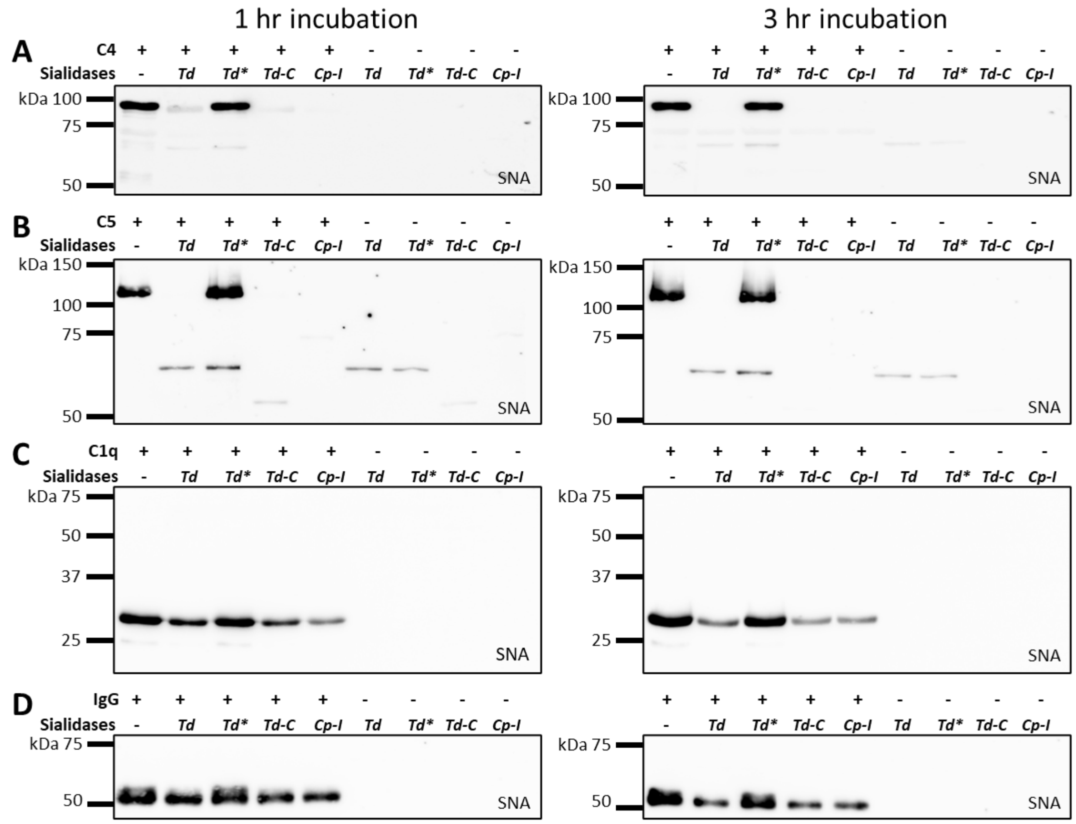

**Figure S10. *Td*-NanH desialylates complement proteins C4 (A), C5 (B), C1q (C), and IgG (D).** For this study, three complement proteins were incubated with sialidases at 37°C for 1 or 3 hours and then subjected to SDS-PAGE followed by lectin blots using SNA lectin. *Td,* the wild type *T. denticola* NanH*; Td**, an catalytic inactive mutant of *Td*-NanH (R140AP142A); *Td-C*; the C-terminal fragment of *Td*-NanH; and *Cp-I*, *C. perfringens* NanI.

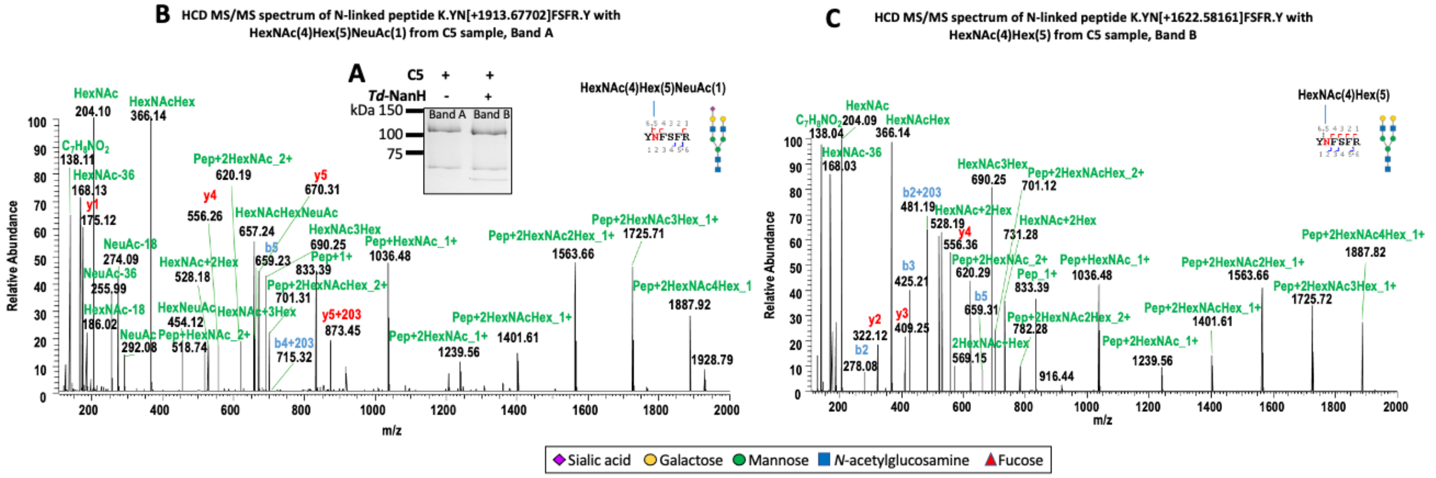

**Figure S11. *Td*-NanH desialylates C5 protein**. (A) SDS-PAGE, followed by Coomassie staining . Untreated (Band A) and treated (Band B) with *Td*-NanH for 3 hours at 37 °C were excised for nano LC-ESI/MS/MS to analyze N-linked glycosylation. **(B,C)** Representative nano LC-ESI-MS/MS spectra of C5 glycoforms from Band A (B) and Band B (C) samples.

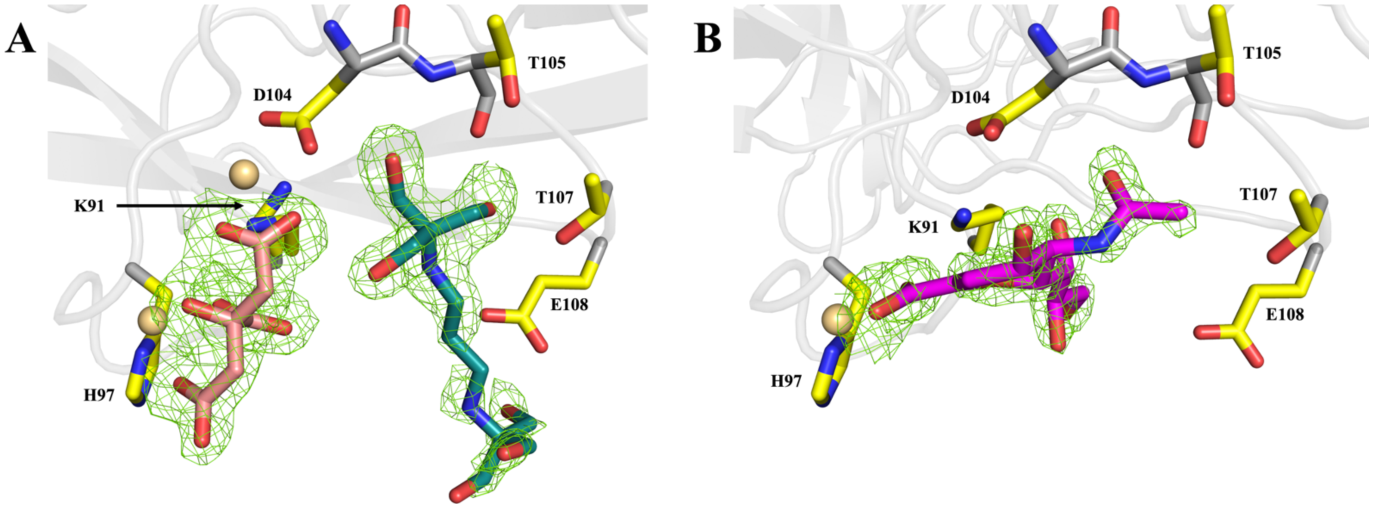


**Figure S12. N0471 Ligand Interactions.** Cartoon representation of the putative ligand-binding site of N0471 bound to **(A)** citrate, bis-tris propane, and cadmium ions and **(B)** DANA. $F_o$-$F_c$ omit electron density maps, contoured at 3$\boldsymbol{\sigma}$, are shown as green mesh. Citrate and bis-tris propane are colored salmon and deep teal for carbons, while nitrogen and oxygen atoms are colored blue and red, respectively. Cadmium ions are shown as wheat spheres. DANA is shown as magenta sticks. Sidechains making interactions with the ligand are colored as yellow, red, and blue for carbons, oxygen, and nitrogen, respectively. Main chain carbons, nitrogen, and oxygen are colored in grey, blue, and red, respectively.

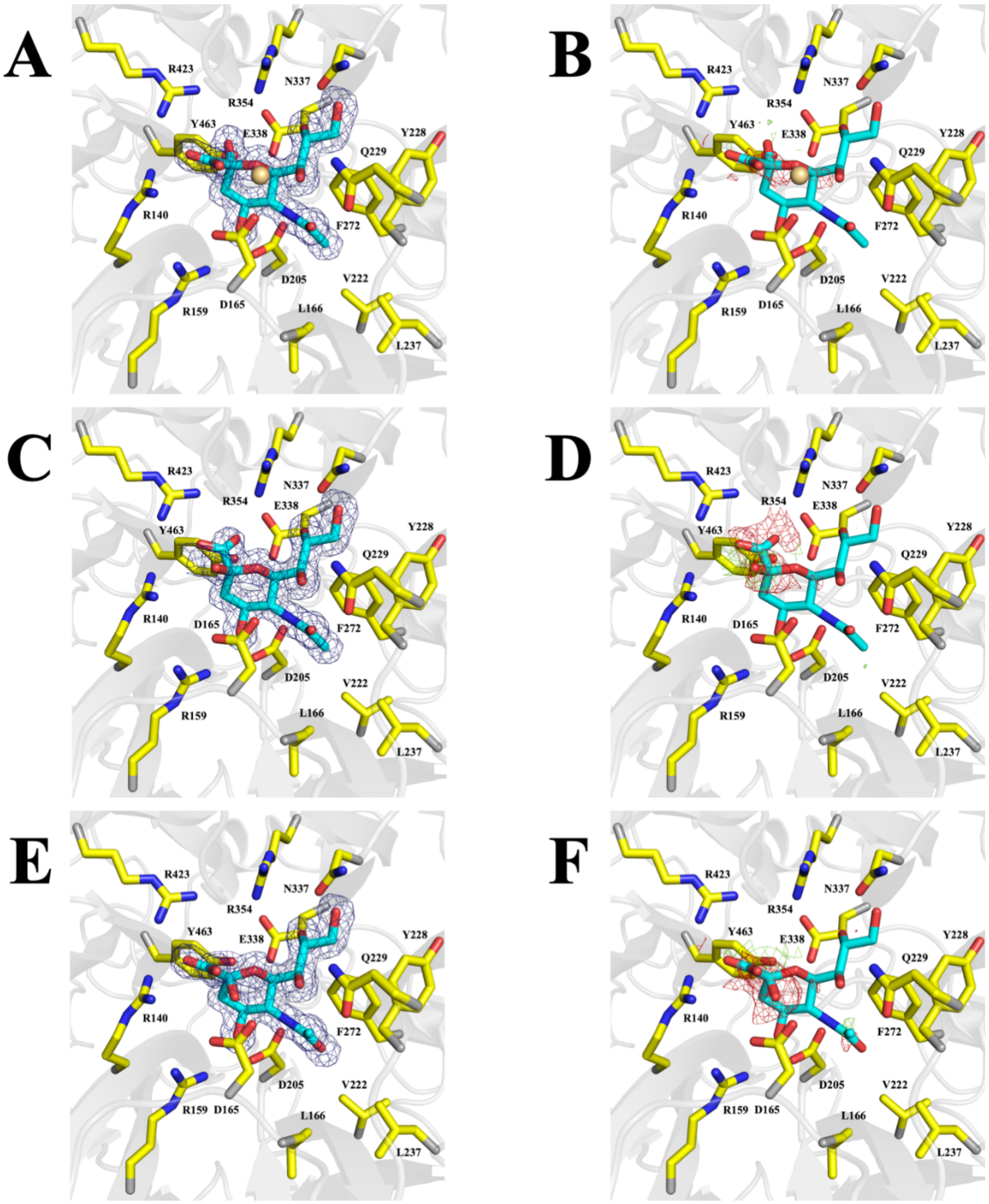


**Figure S13. Validation of β-Neu5Ac modeling.** Cartoon representation of Neu5Ac in complex with TDE0471 modeled as **(A-B)** β-Neu5Ac and **(C-F)** α-Neu5Ac modeled in two different orientations, based on fitting to electron density near the tri-Arg motif. **(A, C, E)** $2mF_o$-$DF_c$ electron density, contoured at 1$\sigma$, and **(B, D, F)** $mF_o$-$DF_c$ difference density, contoured at 3$\sigma$, is shown as blue and green (positive difference density) or red (negative difference density) mesh, respectively. Only the β-anomer **(A/B)** of Neu5Ac fits well to the observed electron and difference density observed. Active site residue side chains are shown as sticks and colored yellow, blue, and red for carbons, nitrogen, and oxygen, respectively. Neu5Ac is shown as sticks and colored teal, blue, and red for carbons, nitrogen, and oxygen, respectively.

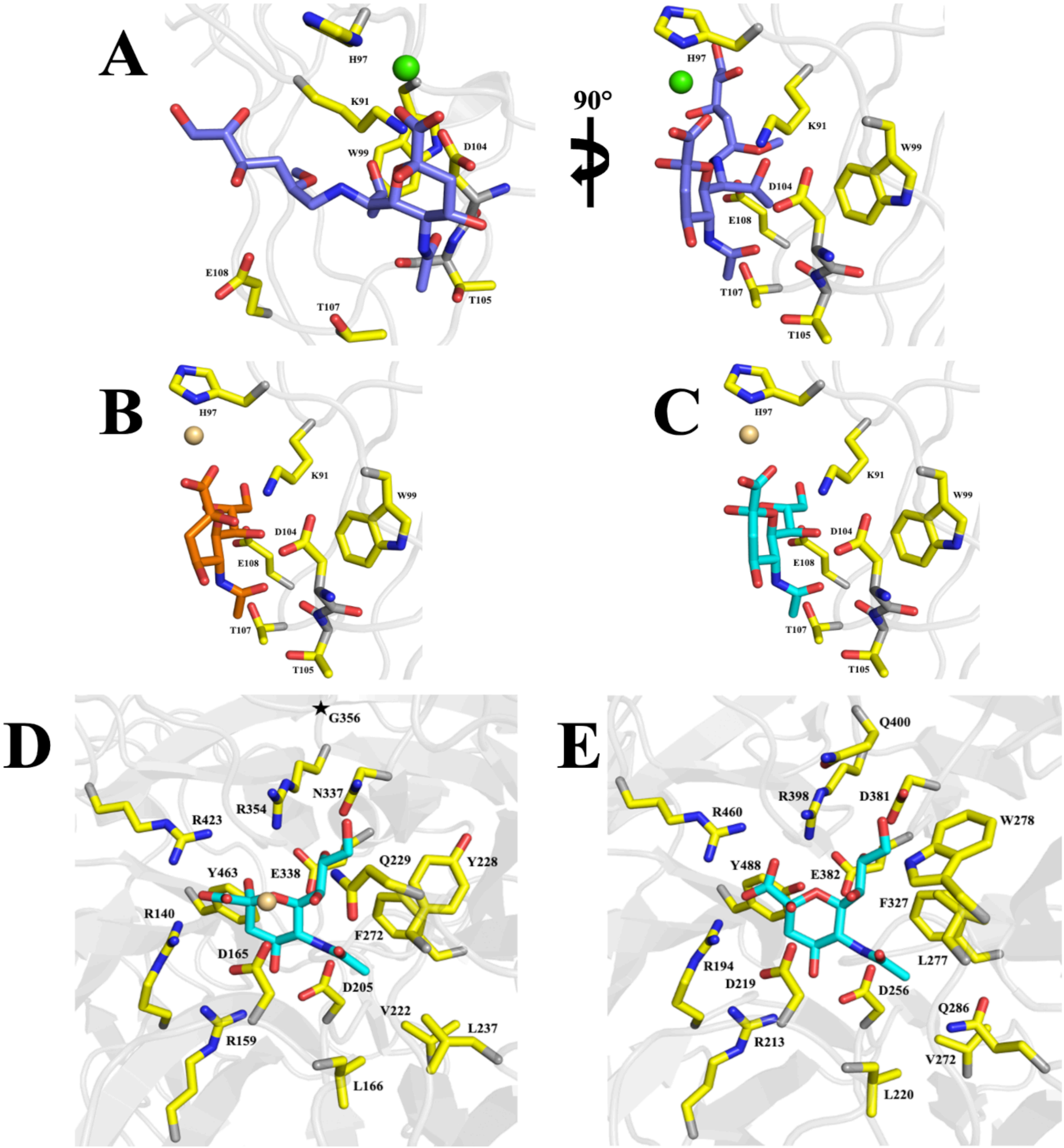


**Figure S14. Assessment of the functional relevance of β-Neu5Ac binding in *Td*-NanH. (A-C)** Cartoon representation of *Td*-NanH:DANA structure, replacing DANA bound in N0471 with the: **(A)** β-anomer of 2,4-dihydroxy-6-[2-hydroxy-1-[[4,5,6-trihydroxy-2-methoxyhexyl]amino]propyl]-5-(1-oxoethylamino)oxane-2-carboxylic acid (Td-SIA; Slate sticks). **(B)** α-anomer of Neu5Ac (orange sticks) **(C)** β-anomer of Neu5Ac (cyan sticks). Calcium is shown as a green sphere, while cadmium is shown as a wheat sphere. **(D-E)** Comparison of active site architecture of **(D)** *Td*-NanH and **(E)** *Porphyromonas gingivalis* sialidase (PDB entry 8FEB). In **(D)** the star icon represents the location of Gly-356, topologically equivalent to Gln-400 in **(E)**. **(A-E)** Sidechains interacting with the modeled substrates are shown as yellow sticks, while mainchain carbons are colored grey, with oxygens and nitrogen colored red and blue, respectively.

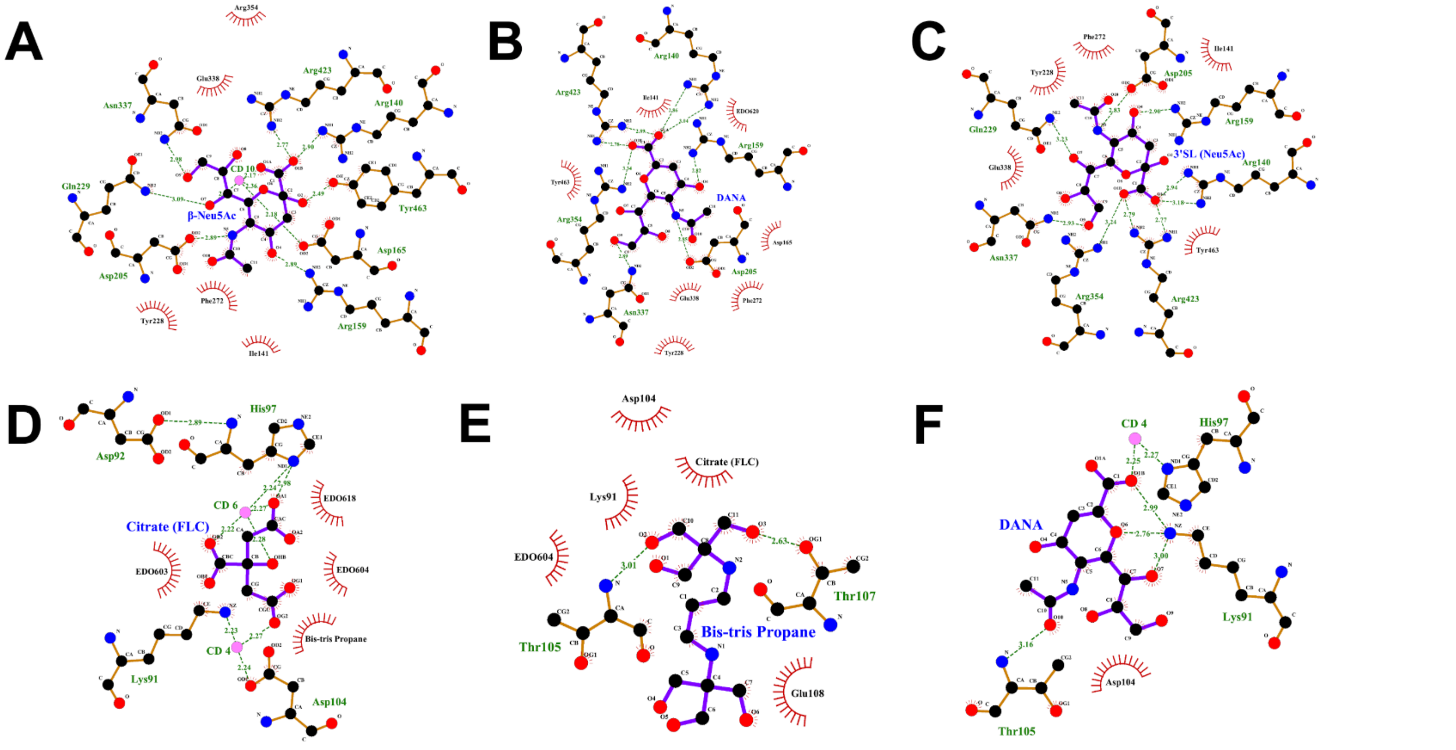


**Figure S15. *LIGPLOT*s Corresponding to *Td*-NanH Ligands.** Depicted are *LIGPLOT*s highlighting the interactions between *Td*-NanH and **(A)** β-Neu5Ac, **(B)** DANA (active site-bound), **(C)** 3'SL **(D)** Citrate, **(E)** Bis-tris propane, and **(F)** DANA (N0471-bound). Note: **(C)** was crystallized with a *Td*-NanH D165A mutant and water molecules were excluded from the *LIGPLOT* analysis.

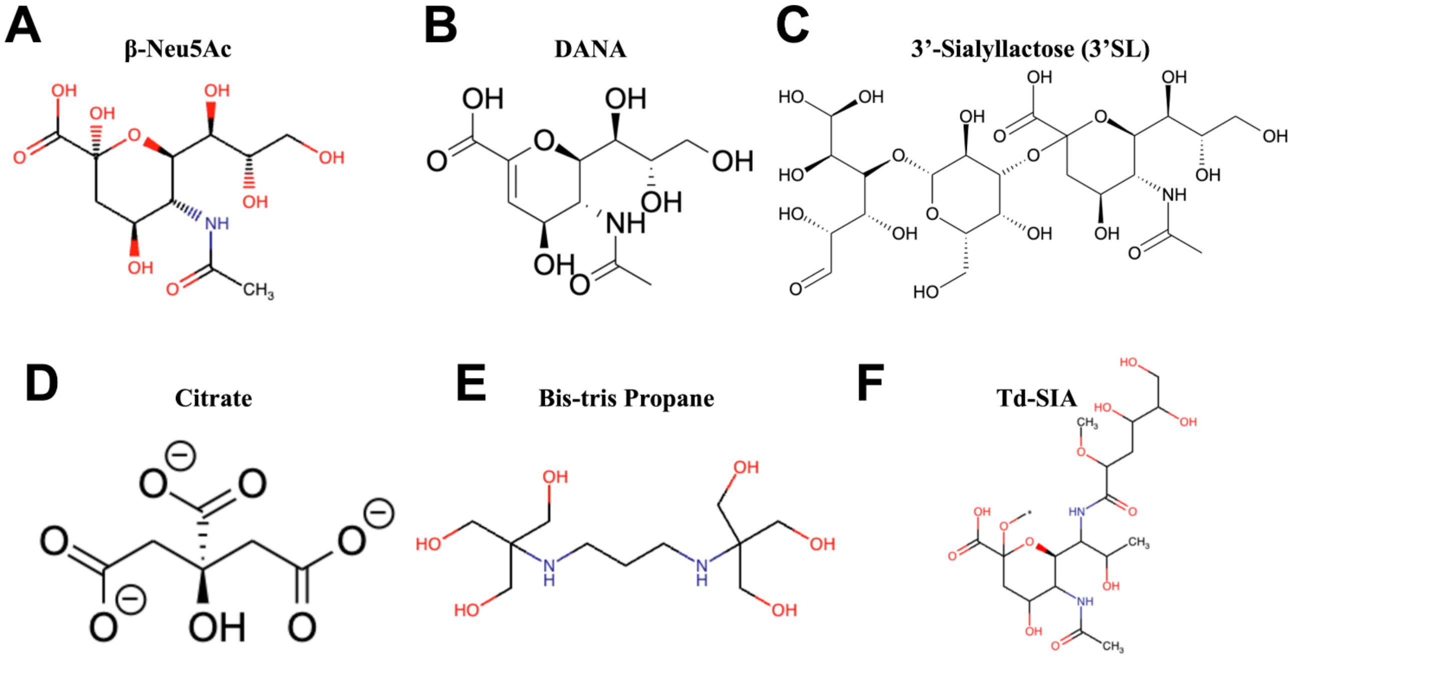


**Figure S16. Chemical structures for ligands used in this study.** Note: for TDE0471:3'SL only density corresponding to the sialic acid (Neu5Ac) moiety was observed (i.e., no density was observed for the galactose and glucose moieties). Note: The * icon in panel F corresponds to where Ser or Thr would be *O*-linked to Td-SIA.

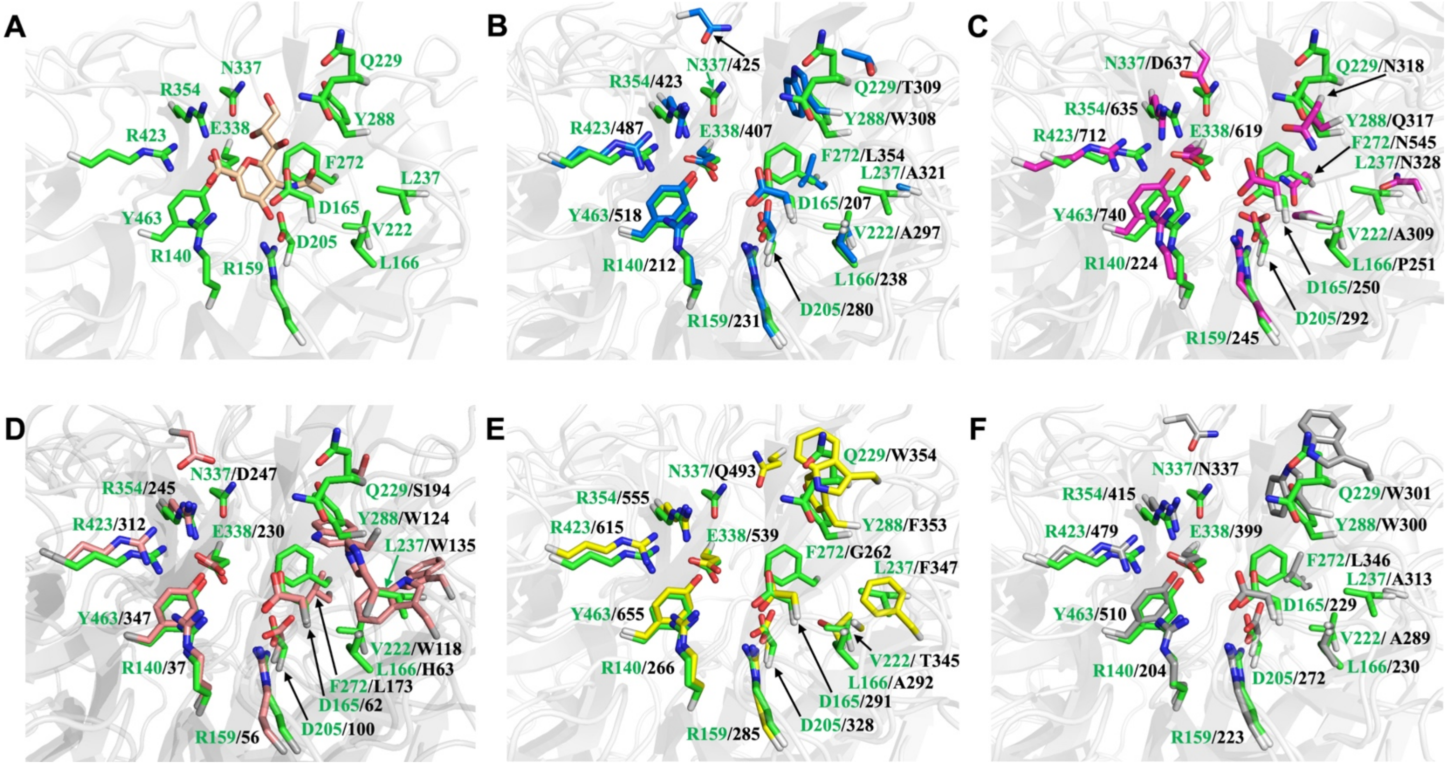


**Figure S17. Active site alignment of six sialidases.** (A-F) Depicted are the active sites of the tested sialidases. Secondary structure is depicted as a grey, transparent cartoon. Side chains depicted as sticks with carbons in green (*Td*-NanH), marine (*Tf*-NanH PDB, 7QYP), magenta (*Vc*-NanH, PDB 1KIT), salmon (*Cp*-NanH, PDB 8UB5), yellow (*Cp*-NanI, PDB 5TSP), and grey (*Bf*-NanH2, AlphaFold) and nitrogen and carbons depicted in blue and red. Residues of *Td*-NanH are labeled in green, while residues of the aligned sialidase is labeled in black. **(A)** Depicts the active site of *Td*-NanH with DANA shown in wheat sticks. **(B-F)** Active site alignment between *Td*-NanH and (B) *Tf*-NanH, (C) *Vc*-NanH, (D) *Cp*-NanH, (E) *Cp*-NanI, and (F) *Bf*-NanH2. Residues near where the carboxylate head of Neu5Ac putatively bind are conserved, while residues interacting with the N-acetyl and glycerol moieties of Neu5Ac are less conserved.
